# Genetic analysis of over one million people identifies 535 novel loci for blood pressure

**DOI:** 10.1101/198234

**Authors:** Evangelos Evangelou, Helen R Warren, David Mosen-Ansorena, Borbala Mifsud, Raha Pazoki, He Gao, Georgios Ntritsos, Niki Dimou, Claudia P Cabrera, Ibrahim Karaman, Fu Liang Ng, Marina Evangelou, Katarzyna Witkowska, Evan Tzanis, Jacklyn N Hellwege, Ayush Giri, Digna R Velez Edwards, Yan V Sun, Kelly Cho, J.Michael Gaziano, Peter WF Wilson, Philip S Tsao, Csaba P Kovesdy, Tonu Esko, Reedik Mägi, Lili Milani, Peter Almgren, Thibaud Boutin, Stéphanie Debette, Jun Ding, Franco Giulianini, Elizabeth G Holliday, Anne U Jackson, Ruifang Li-Gao, Wei-Yu Lin, Jian'an Luan, Massimo Mangino, Christopher Oldmeadow, Bram Prins, Yong Qian, Muralidharan Sargurupremraj, Nabi Shah, Praveen Surendran, Sébastien Thériault, Niek Verweij, Sara M Willems, Jing-Hua Zhao, Philippe Amouyel, John Connell, Renée de Mutsert, Alex SF Doney, Martin Farrall, Cristina Menni, Andrew D Morris, Raymond Noordam, Guillaume Paré, Neil R Poulter, Denis C Shields, Alice Stanton, Simon Thom, Gonçalo Abecasis, Najaf Amin, Dan E Arking, Kristin L Ayers, Caterina M Barbieri, Chiara Batini, Joshua C Bis, Tineka Blake, Murielle Bochud, Michael Boehnke, Eric Boerwinkle, Dorret I Boomsma, Erwin P Bottinger, Peter S Braund, Marco Brumat, Archie Campbell, Harry Campbell, Aravinda Chakravarti, John C Chambers, Ganesh Chauhan, Marina Ciullo, Massimiliano Cocca, Francis Collins, Heather J Cordell, Gail Davies, Martin H de Borst, Eco J de Geus, Ian J Deary, Joris Deelen, Fabiola M Del Greco, Cumhur Yusuf Demirkale, Marcus Dörr, Georg B Ehret, Roberto Elosua, Stefan Enroth, A.Mesut Erzurumluoglu, Teresa Ferreira, Mattias Frånberg, Oscar H Franco, Ilaria Gandin, Paolo Gasparini, Vilmantas Giedraitis, Christian Gieger, Giorgia Girotto, Anuj Goel, Alan J Gow, Vilmundur Gudnason, Xiuqing Guo, Ulf Gyllensten, Anders Hamsten, Tamara B Harris, Sarah E Harris, Catharina A Hartman, Aki S Havulinna, Andrew A Hicks, Edith Hofer, Albert Hofman, Jouke-Jan Hottenga, Jennifer E Huffman, Shih-Jen Hwang, Erik Ingelsson, Alan James, Rick Jansen, Marjo-Riitta Jarvelin, Roby Joehanes, Åsa Johansson, Andrew D Johnson, Peter K Joshi, Pekka Jousilahti, J Wouter Jukema, Antti Jula, Mika Kähönen, Sekar Kathiresan, Bernard D Keavney, Kay-Tee Khaw, Paul Knekt, Joanne Knight, Ivana Kolcic, Jaspal S Kooner, Seppo Koskinen, Kati Kristiansson, Zoltan Kutalik, Maris Laan, Marty Larson, Lenore J Launer, Benjamin C Lehne, Terho Lehtimäki, Daniel Levy, David CM Liewald, Li Lin, Lars Lind, Cecilia M Lindgren, Yongmei Liu, Ruth JF Loos, Lorna M Lopez, Lingchan Lu, Leo-Pekka Lyytikäinen, Anubha Mahajan, Chrysovalanto Mamasoula, Jaume Marrugat, Jonathan Marten, Yuri Milaneschi, Anna Morgan, Andrew P Morris, Alanna C Morrison, Peter J Munson, Mike A Nalls, Priyanka Nandakumar, Christopher P Nelson, Christopher Newton-Cheh, Teemu Niiranen, Ilja M Nolte, Teresa Nutile, Albertine J Oldehinkel, Ben A Oostra, Paul F O'Reilly, Elin Org, Sandosh Padmanabhan, Walter Palmas, Arno Palotie, Alison Pattie, Brenda WJH Penninx, Markus Perola, Annette Peters, Ozren Polasek, Peter P Pramstaller, Nguyen Quang Tri, Olli T Raitakari, Meixia Ren, Rainer Rettig, Kenneth Rice, Paul M Ridker, Janina S Reid, Harriëtte Riese, Samuli Ripatti, Antonietta Robino, Lynda M Rose, Jerome I Rotter, Igor Rudan, Daniella Ruggiero, Yasaman Saba, Cinzia F Sala, Veikko Salomaa, Nilesh J Samani, Antti-Pekka Sarin, Rheinhold Schmidt, Helena Schmidt, Nick Shrine, David Siscovick, Albert V Smith, Harold Schneider, Siim Sõber, Rossella Sorice, John M Starr, David J Stott, David P Strachan, Rona J Strawbridge, Johan Sundström, Morris A Swertz, Kent D Taylor, Alexander Teumer, Martin D Tobin, Daniela Toniolo, Michela Traglia, Stella Trompet, Jaakko Tuomilehto, Christophe Tzourio, André G Uitterlinden, Ahmad Vaez, Peter J van der Most, Cornelia M van Duijn, Anne-Claire Vergnaud, Germaine C Verwoert, Veronique Vitart, Uwe Völker, Peter Vollenweider, Dragana Vuckovic, Hugh Watkins, Sarah H Wild, Gonneke Willemsen, James F Wilson, Alan F Wright, Jie Yao, Tatijana Zemunik, Weihua Zhang, John R Attia, Adam S Butterworth, Dan Chasman, David Conen, Francesco Cucca, John Danesh, Caroline Hayward, Joanna MM Howson, Markku Laakso, Edward G Lakatta, Claudia Langenberg, Ollie Melander, Dennis O Mook-Kanamori, Patricia B Munroe, Colin NA Palmer, Lorenz Risch, Robert A Scott, Rodney J Scott, Peter Sever, Tim D Spector, Pim van der Harst, Nicholas J Wareham, Eleftheria Zeggini, Morris J Brown, Andres Metspalu, Adriana M Hung, Christopher J O'Donnell, Todd L Edwards, on behalf of the Million Veteran Program, Bruce M. Psaty, Ioanna Tzoulaki, Michael R Barnes, Louise V Wain, Paul Elliott, Mark J Caulfield

## Abstract

High blood pressure is the foremost heritable global risk factor for cardiovascular disease. We report the largest genetic association study of blood pressure traits to date (systolic, diastolic, pulse pressure) in over one million people of European ancestry. We identify 535 novel blood pressure loci that not only offer new biological insights into blood pressure regulation but also reveal shared loci influencing lifestyle exposures. Our findings offer the potential for a precision medicine strategy for future cardiovascular disease prevention.

---

High blood pressure, defined as a systolic blood pressure >140 mm Hg and/or diastolic blood pressure > 90 mm Hg, is the dominant inherited risk factor for stroke and coronary artery disease and is responsible for an estimated 7.8 million deaths and 148 million disability life years lost worldwide in 2015 alone^1^. Family studies indicate that the blood pressure (BP) level is determined in an individual by complex interactions between life course exposures and their genetic background^2–5^. Previous genetic association studies have included genome-wide meta-analyses, customised cardiovascular candidate centric analyses and evaluation of exome variation that have identified and validated variants at 274 loci which have a modest effect on population BP and explain ~3-4% of the trait variance^6–13^.

Here, we report genome-wide discovery analyses of BP traits (systolic - SBP, diastolic - DBP and pulse pressure-PP) in people of European ancestry drawn from UK Biobank (UKB)^14^ and the International Consortium of Blood Pressure-Genome Wide Association Studies (ICBP)^12,13^. We adopted a combination of a one and two-stage study design to test common and low-frequency single nucleotide polymorphisms (SNPs) with minor allele frequency (MAF) ≥ 1% in association with BP traits (**Fig. 1**) including over 1 million people of European descent across both discovery and replication, including data from the Million Veterans Program (MVP) and the Estonian Biobank.

**Figure 1.**
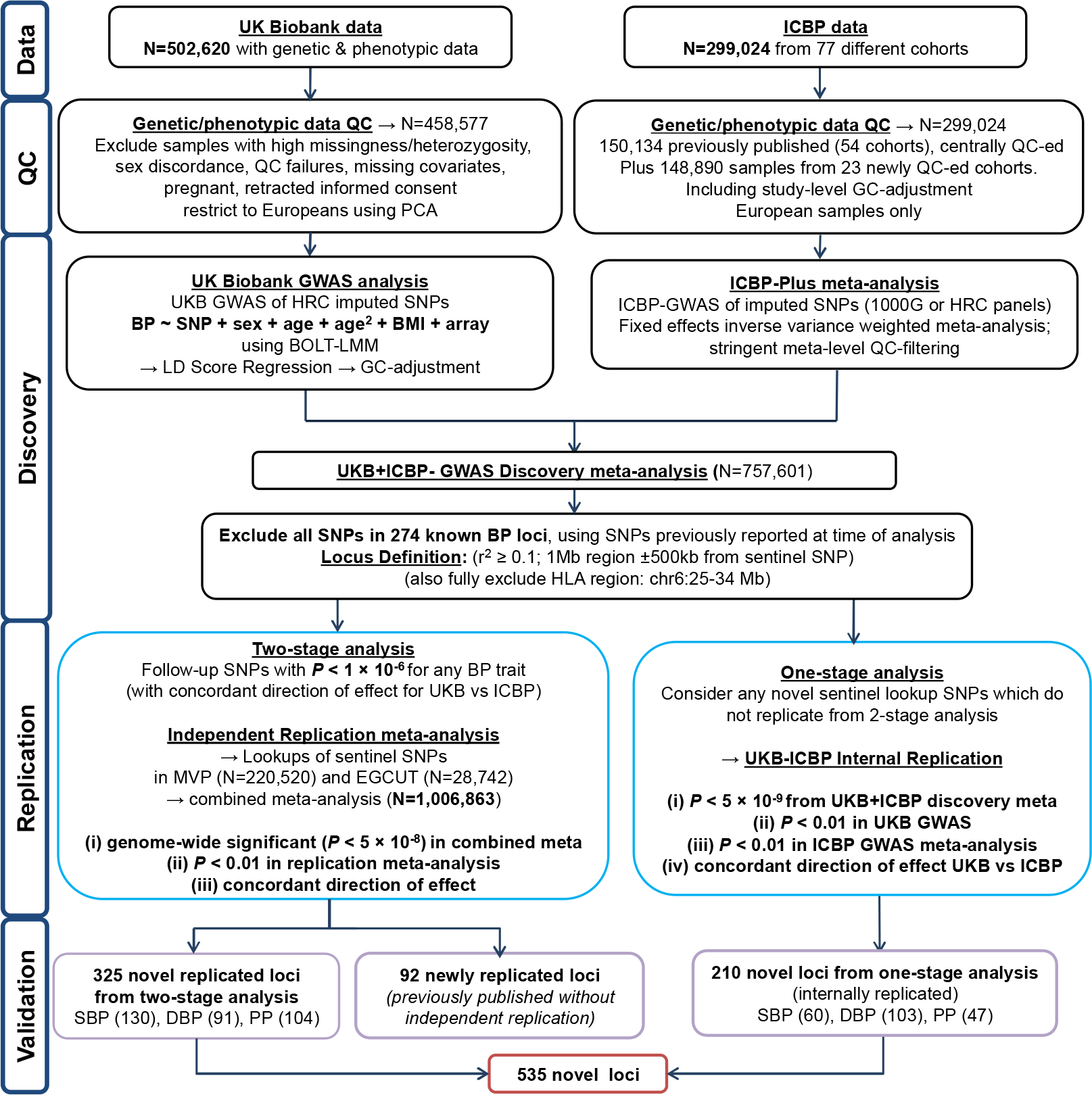
Study design. Schematic of the one- and two-stage design. ICBP; International Consortium for Blood Pressure; N, sample size; QC, quality control; PCA, principal-component analysis; GWAS, Genome-wide Association Study; 1000G 1000 Genomes; HRC, Haplotype Reference Panel; BP: blood pressure; SNPs, single nucleotide polymorphisms; BMI, body mass index; LMM; linear mixed model; UKB, UK Biobank, MAF, minor allele frequency; HLA, Human Leukocyte Antigen; MVP, Million Veterans Program; EGCUT; Estonian Genome Center, University of Tartu; SBP, systolic blood pressure; DBP, diastolic blood pressure; PP, pulse pressure.

Briefly, UKB is a prospective cohort study of ~500,000 individuals recruited at ages 40-69 years who have been richly phenotyped including BP measurements^14^. Participants were genotyped using a customized array with imputation from the Haplotype Reference Consortium (HRC) panel, yielding ~7 million SNPs (imputation quality score (INFO) ≥ 0. 1)^15^. After quality control (QC) and exclusions we performed genome-wide association studies (GWAS) of BP traits using data from 458,577 individuals of European descent under an additive genetic model^16^ (**Supplementary Table 1a**). Following LD-score regression^17^, genomic control was applied to the UKB data prior to meta-analysis.

In addition, we performed GWAS analyses for BP traits in the ICBP GWAS data comprising 77 independent studies including up to 299,024 participants of European ancestry genotyped with various arrays, and imputed to either 1,000 Genomes or the HRC platforms (**Supplementary Table 1b**). After QC we applied genomic control at study-level and obtained summary effect sizes on ~7 million SNPs with INFO ≥ 0.3 and Cochran’s Q statistic ^18^ (test of heterogeneity) filtered at *P* ≥ 1 × 10^−4^.

We then combined the UKB and ICBP GWAS results using inverse-variance weighted fixed effects meta-analysis, giving a total discovery sample of 757,601 individuals^19^.

In our two-stage design we attempted replication of 1,062 SNPs at *P* < 1 × 10^−6^ from discovery with concordant effect direction between UKB and ICBP, using the sentinel SNP (i.e. SNP with smallest *P*-value at the locus) after excluding the HLA region (chr 6:25-34MB) and all SNPs in Linkage Disequilibrium (LD) (r^2^ ≥ 0.1) or ±500 Kb from any previously validated BP-associated SNPs at the 274 published loci. We used the US Million Veteran Program (MVP, up to 220,520 people of European descent) and the Estonian Genome Centre Biobank (EGCUT, up to 28,742 Europeans) as our replication resources^20,21^ (**Supplementary Table 1c**). Our replication criteria for the two-stage design were genome-wide significance (*P* < 5 × 10^−8^) in the combined meta-analysis, with *P* < 0.01 in the replication data and concordant direction of effect between discovery and replication.

Given the large size of the two discovery resources (UKB and ICBP) we additionally undertook a one-stage design with internal replication, to minimize the risk of missing true positive associations from our two-stage analysis. To ensure the robustness of this approach, and to avoid false positive findings, we used *P* < 5 × 10^−9^ as the *P*-value threshold from the discovery meta-analysis, a criterion that is an order of magnitude more stringent than genome-wide significance^22^. We also required an internal replication *P*-value of < 0.01 in each of the UKB and ICBP GWAS analyses and with concordant direction of effect.

We then explored the putative function of the BP associated signals using a range of *in silico* resources, including expression quantitative trait loci (eQTLs), tissue and DNase I site enrichment, long range chromatin interactions (Hi-C), pathway analysis and druggability. We investigated metabolomic signatures associated with our novel sentinel SNPs, evaluated the overlap with lifestyle exposures that influence BP, and examined the overlap of BP-associated loci with other complex traits and diseases. Finally, we performed a genetic risk score analysis to model the impact that all BP-associated variants have on BP level, risk of hypertension (HTN), other cardiovascular diseases and on BP in other non-European ancestries.

## RESULTS

We report a total of 535 novel loci, of which 325 were identified from our two-stage design and a further 210 from our one-stage design with internal replication (**Fig. 2**, **Fig. 3a** and **Supplementary Tables 2a-2f**). We note that the two designs identified large numbers of distinctive loci as well as some overlap, with similar distribution of effect sizes, illustrating the power and complementarity of our dual approach (Fig. 3a and **Supplementary Fig. 2**). In addition, we confirmed a further 92 loci that had been previously reported but not replicated (**Supplementary Table 3**)^10^. We also confirm previous findings for all 274 published BP loci (**Supplementary Fig. 1 & 2** and **Supplementary Table 4**). Our findings bring the total number of BP loci to 901, more than tripling the number of previously published BP loci. Similarly, there is substantial gain in the percentage of BP explained, for instance increasing the variance explained by ~2.5-fold for SBP from 4.6 % for the 274 published loci to 11.2% for all 901 loci.

**Figure 2.**
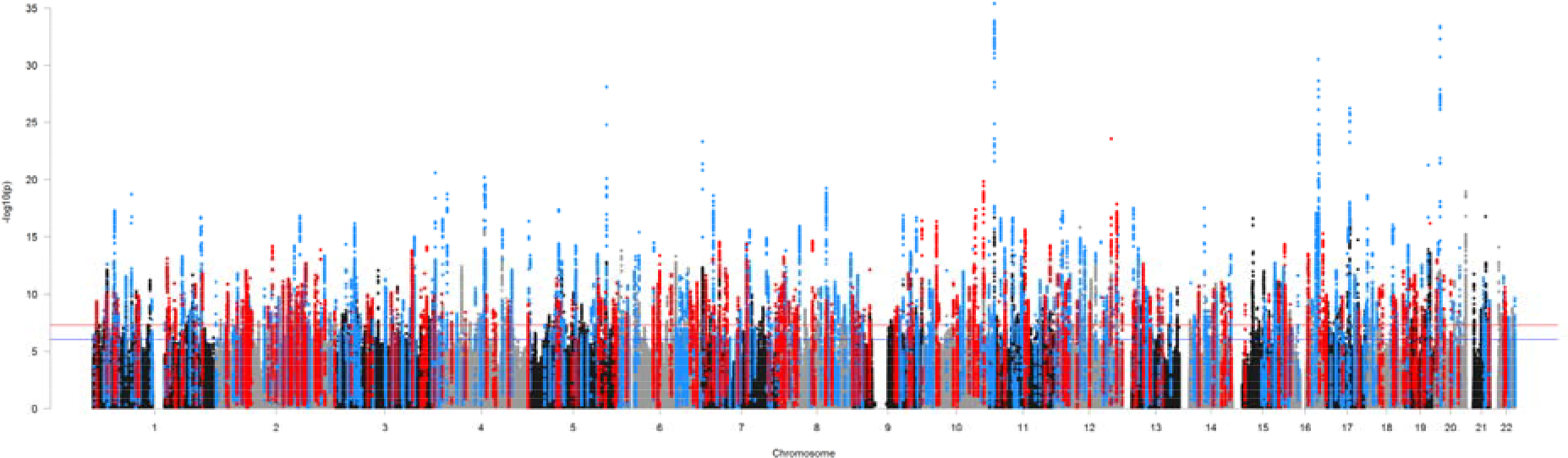
Manhattan plot showing the minimum *P*-value for the association across all blood pressure traits in the discovery stage excluding previously published variants. Manhattan plot of the discovery genome-wide association meta-analysis in >750,000 individuals excluding variants in published loci. The minimum *P*-value across SBP, DBP and PP is presented. The y-axis shows the – log_10_ *P* values and the x- axis shows their chromosomal positions. Horizontal red and blue line represents the thresholds of *P* = 5 × 10^−8^ for genome-wide significance and *P* = 1 × 10^−6^ for selecting SNPs for replication, respectively. SNPs in blue are in LD (r^2^ > 0.8) with the 325 novel variants from the two-stage design whereas SNPs in red are in LD (r^2^ > 0.8) with 210 SNPs identified through the one-stage design with internal replication. Any loci in black or grey that exceed the significance thresholds were significant in the discovery meta-analysis, but did not meet the criteria of replication in the one - or two-stage designs.

**Figure 3:**
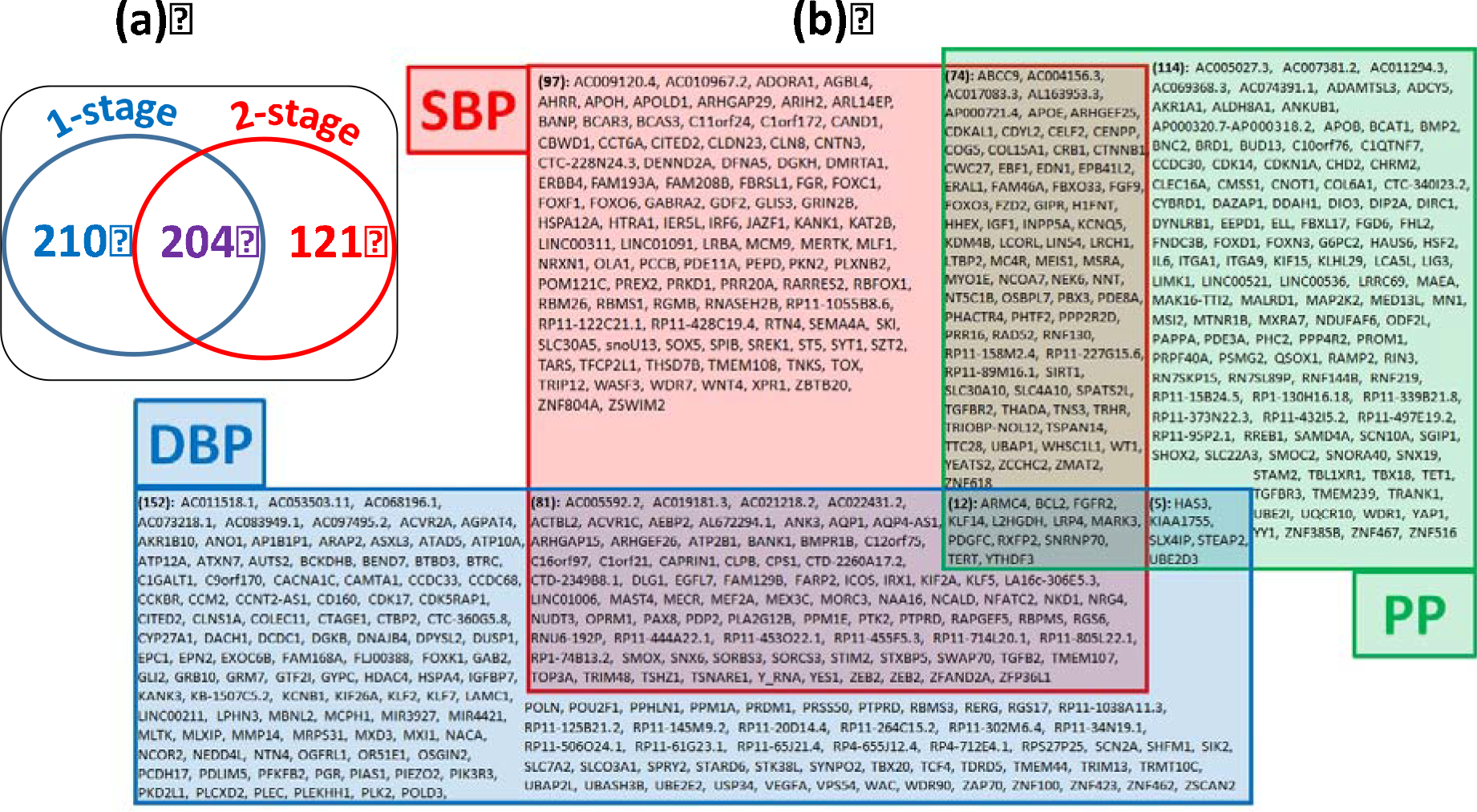
Venn Diagrams of Novel Loci Results (a) “Comparison of one-stage and two-stage design analysis criteria”: For all 535 novel loci, we compare the results according to the association criteria used between the one-stage design analysis and the two-stage design analysis. The 210 novel loci from the one-stage analysis were selected from the remaining lookup SNPs which did not replicate from the 2-stage analysis (i.e. did not reach genome-wide significance with *P* < 0.01 in the MVP+EGCUT replication and concordant effect direction for any traits). But we show that, of the 325 novel replicated loci from the 2-stage analysis, 204 loci would also have met the one-stage criteria (with *P* <5×10^−9^ in the discovery meta-analysis, *P* < 0.01 in UKB, *P* < 0.01 in ICBP and concordant direction of effect between UKB and ICBP), whereas 121 would only have been identified from the two-stage analysis. **(b) “Overlap of Associations across Blood Pressure Traits”**: For all 535 novel loci, we show which blood pressure traits each locus was associated with. For the 325 novel replicated loci from two-stage analysis, we show the traits that met the two-stage analysis criteria with genome-wide significance in the combined meta-analysis, *P* < 0.01 significance in the replication meta-analysis and concordant direction of effect. For the 210 novel loci from one-stage analysis, we show the traits that met the one-stage analysis criteria with *P* < 5×10^−9^ in the discovery meta-analysis, *P* < 0.01 in UKB, *P* < 0.01 in ICBP and concordant direction of effect between UKB and ICBP. The locus names provided in alphabetical order correspond to the nearest annotated gene. SNPs: Single nucleotide polymorphisms; SBP: systolic blood pressure; DBP: diastolic blood pressure; PP: pulse pressure; UKB: UK Biobank; ICBP: International Consortium of Blood Pressure; MVP: Million Veterans Program; EGCUT: Estonian Genome Center, University of Tartu.

### Discovery of novel genetic loci for blood pressure

Of the 535 independent novel loci, 363 were associated with only one trait; 160 with two traits and 12 loci with all three BP traits (**Fig. 3**), reflecting the inter-correlations between BP traits despite their different physiology (**Supplementary Fig. 2**). The largest number of novel loci per primary trait (the most significantly associated trait from the one- or two-stage analyses) was for DBP (194 loci), followed by SBP (190) and PP (151) (**Supplementary Table 2a-f**).

### Functional analyses

Our functional analyses approach is summarised in **Supplementary Figure 3**. First, we annotated all 901 loci to 1644 novel and 962 known genes (based on LD r^2^ ≥ 0.8) and then investigated these loci for tissue enrichment, DNase hypersensitivity site enrichment and pathway analyses. At 66 of the 535 novel loci we identified 97 non-synonymous SNPs, including 8 predicted to be damaging (**Supplementary Table 5**).

We used chromatin interaction Hi-C data from endothelial cells (HUVEC)^23^, neural progenitor cells (NPC), mesenchymal stem cells (MSC) and tissue from the aorta and adrenal gland^24^ to identify distal affected genes. There were 498 novel loci that contained a potential regulatory SNP and in 484 we identified long-range interactions in at least one of the tissues or cell types. We found several potential long-range target genes that do not overlap with the sentinel SNPs in the LD block, such as *TGFB2* that forms a 1.2Mb long regulatory loop with the SNPs in the *SLC30A10* locus, and the *TGFBR1* promoter that forms a 100kb loop with the *COL15A1* locus **(Supplementary Table 5)**.

Our eQTL analysis identified 92 novel loci with eQTLs in arterial tissue and 48 in adrenal tissue **(Supplementary Table 6);** these are more compared to what was identified in our previously published UKB GWAS^11^. An example is the rs31120122 SNP that defines an aortic eQTL that affects expression of the *MED8* gene within the *SZT2* locus. In combination with Hi-C interaction data in MSC this finding supports a role *MED8* in BP regulation, possibly mediated through repression of smooth muscle cell differentiation. Hi-C interactions provide confirmatory evidence for involvement of a further 13 arterial eGenes (genes whose expression is affected by the eQTLs) that were distal to their eQTLs (*MECR*, *CCDC30*, *DNAJC9-AS1*, *DCAF16*, *POU4F1*, *ANKDD1A*, *EPN2*, *MRPS6*, *CDYL2*, *WDR92*, *LINC00310*, *TAPT1*, *GBR10*).

We investigated which transcription factors and chromatin marks are involved in regulatory interactions using the functional predictions from DeepSEA. We found 198 SNPs in 121 novel loci with predicted effects on transcription factor binding or on chromatin marks in tissues relevant for BP biology, such as vascular tissue, smooth muscle and the kidney **(Supplementary Table 5)**.

We used our genome wide data at a false discovery rate (FDR) < 1% to robustly assess the tissue enrichment of BP loci using DEPICT and identified enrichment across 50 tissues and cells (**Supplementary Fig 4**; **Supplementary Table 7a**). Enrichment was greatest for the cardiovascular system especially blood vessels (*P* = 1.5 × 10^−11^) and the heart (*P* = 2.7 × 10^−5^). Enrichment was high in adrenal tissue (*P* = 3.7 × 10^−4^) and, for the first time, we observed high enrichment in adipose tissues (*P* = 9.8 × 10^−9^) corroborated by eQTL enrichment analysis (*P* < 0.05) **(Supplementary Fig. 4; Supplementary Table 7a)**. Evaluation of enriched mouse knockout phenotype terms also points to the importance of vascular morphology (*P* = 6 × 10^−15^) and development (*P* =2.1 × 10^−18^) in BP. Due to the contribution of our novel BP loci, we identify new findings from both the gene ontology and protein-protein interaction subnetwork enrichments, which highlighted the TGFβ (*P* = 2.3 × 10^−13^) and related SMAD pathways (*P* = 7 × 10^−15^) (**Supplementary Table 7b, Supplementary Fig. 5b-d**).

We used FORGE to investigate the regulatory regions for cell type specificity from DNase I hypersensitivity sites, which showed strongest enrichment (*P* < 0.001) in the vasculature and highly vascularised tissues, similar to previous BP studies^11^ **(Supplementary Fig. 6).**

Ingenuity pathway analysis and upstream regulator assessment shows enrichment of canonical pathways implicated in cardiovascular disease including pathways targeted by antihypertensive drugs (e.g. nitric oxide signalling) and also suggests some potential new targets, such as relaxin signalling. Notably, upstream regulator analysis identified several known mediators of BP including some therapeutic targets such as angiotensinogen, calcium channels, progesterone, natriuretic peptide receptor, angiotensin converting enzyme, angiotensin receptors and endothelin receptors **(Supplementary Fig. 7)**.

### Potential therapeutic targets

We present a summary of the vascular expressed genes contained within the 535 novel loci identified, including a review of their potential druggability **(Supplementary Fig. 8)**. The overlap between genes associated with blood pressure and genes associated with antihypertensive drug targets, demonstrates new genetic support for known drug mechanisms. For example, we report novel BP associations at the *PKD2L1* and *BCL2* loci, which are targeted by amiloride and atenolol respectively. Extending associations by LD we report novel genetic associations with the targets of eleven antihypertensive drugs **(Supplementary Table 8).**

### Metabolomic analysis of BP associated variants

We used ^1^H NMR lipidomics data on plasma and data from the Metabolon platform for a subset of 2,022 and 1,941 participants of the Airwave Health Monitoring Study respectively^25^ for our sentinel SNPs. We also used PhenoScanner to test each SNP against published significant (*P* < 5 × 10^−8^) genome vs metabolome-wide associations in plasma and urine.

Ten SNPs show association with lipid particle metabolites and a further 31 SNPs (8 also on PhenoScanner) show association with metabolites on the Metabolon platform. These highlight lipid pathways, amino acids (glycine, serine, glutamine), tri-carboxylic acid cycle intermediates (succinylcarnitine) and drug metabolites (**Supplementary Table 9 and 10**). These findings suggest a close metabolic coupling of BP regulation with lipid, and for the first time, with energy metabolism.

### Concordance of BP Variants and Lifestyle exposure

UK Biobank has collected extensive lifestyle related data which have been epidemiologically associated with BP. These include macronutrient, water, tea, coffee and alcohol intake, anthropomorphic traits, physical activity and inactivity, smoking and urinary sodium, potassium and creatinine excretion^14^. We investigated whether sentinel SNPs at all 901 BP loci were associated with these traits in the Stanford Global Biobank Engine (N = 327,302 urinary traits) and Gene ATLAS (N = 408,455) for all other traits, with corrected *P* < 1 × 10^−6^). We find that a BP SNP rs34783010 in *GIPR* is associated with daily fruit intake (*P* = 1.03 × 10^−7^), urinary sodium and creatinine concentration (*P* = 1.5 × 10^−13^ and 1.2 × 10^−9^ respectively), body mass index (*P* = 3.3 × 10^−41^), weight (*P* = 7.3 × 10^−35^) and waist circumference (*P* = 7.7 × 10^−30^); rs6495122, near *CPLX3* and *ULK3*, and rs1378942, in *CSK* are associated with water (*P* = 1.3 × 10^−22^ and 2.6 × 10^−20^ respectively), caffeine (*P* = 1.3 × 10^−46^ and 2.2 × 10^−43^) and tea intake (*P* = 7.6 × 10^−38^ and 8.1 × 10^−33^), as well as urinary creatinine concentrations (*P*= 5.6 × 10^−8^ and *P*= 3.2 × 10^−8^ respectively). In addition, the BP SNP rs13107325 in *SLC39A8*, is a novel locus for frequency of drinking alcohol (*P* = 3.5 × 10^−15^) and time spent watching TV (*P* = 2.3 × 10^−11^) as well as being associated with body mass index (*P* = 1.6 × 10^−33^), weight (*P* = 8.8 × 10^−16)^ and waist circumference (*P* = 4.7 × 10^−11^) **(Supplementary table 11a).** We used unsupervised hierarchical clustering for the 36 BP loci that showed at least one association with the lifestyle or anthropometric traits in UKB at *P* < 1 × 10^−6^ (**Fig. 4**). The heatmap summarises the locus specific associations across the range of traits and highlights heterogeneous effects with anthropometric traits across the range of loci examined. For example, it shows a cluster of associations between BP raising alleles and increased adult height and weight and another cluster of genes that show associations between BP raising alleles and decreased adult height and weight.

**Figure 4.**
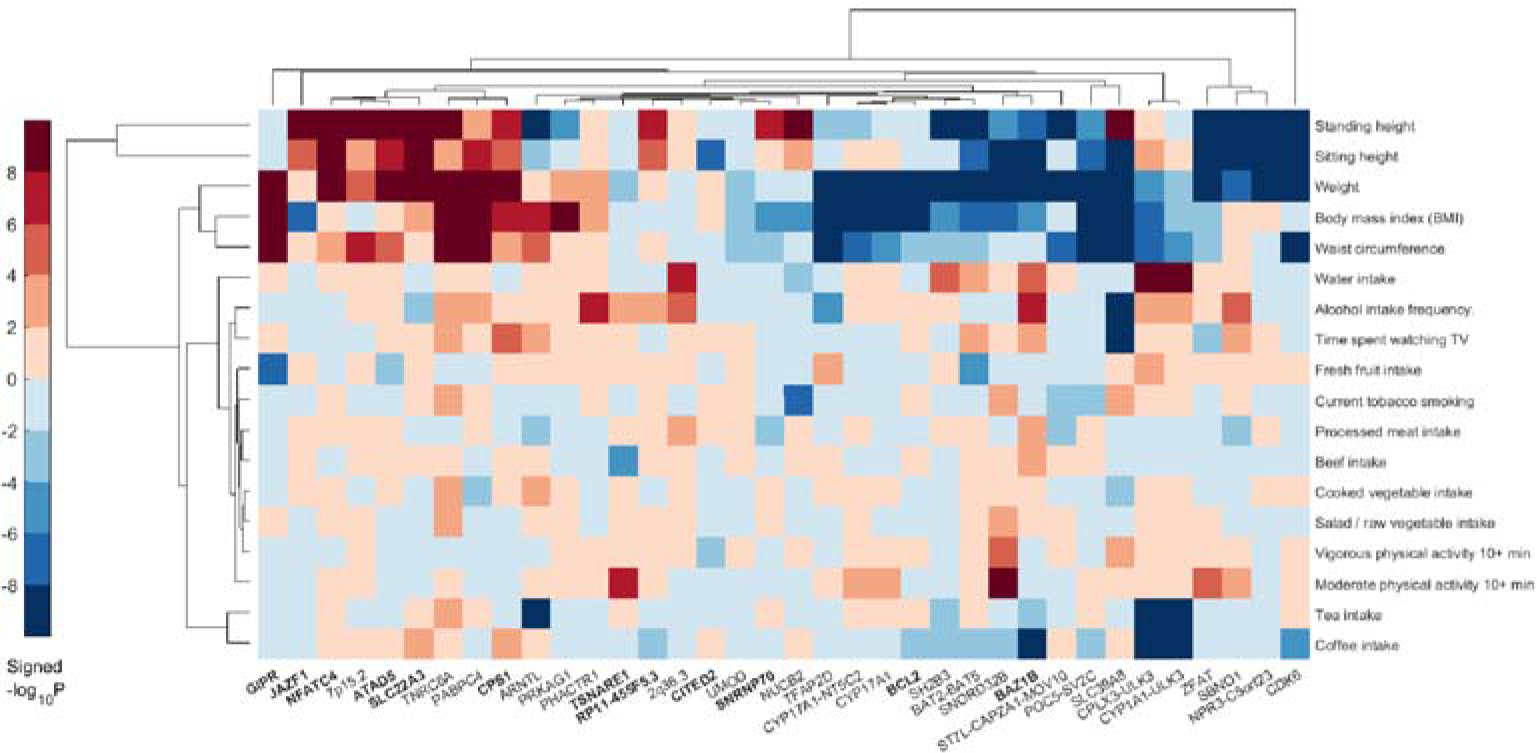
Association of blood pressure loci with lifestyle traits. Plot shows hierarchical clustering of BP loci based on associations with anthropometry and lifestyle factors. For the lead SNP at each BP locus (x-axis), we calculated the −log_10_(*P*)*sign(β) (aligned to BP-raising allele) as retrieved form the Gene Atlas (http://geneatlas.roslin.ed.ac.uk). BP loci and traits were clustered according to the Euclidean distance amongst −log_10_(*P*)*sign(β). Red squares indicate direct associations with the trait of interest and blue squares inverse associations. Only SNPs with at least one association with *P* <1 × 10^−6^ with at least one of the traits examined are annotated in the heat-map. All 901 loci are considered, both published and novel: novel loci are printed in bold font. SNPs: Single Nucleotide Polymorphisms; BP: Blood Pressure.

### Association lookups with other traits and diseases

We evaluated cross-trait effects using PhenoScanner^26^ and DIsGeNET, which integrates data from expert curated repositories, GWAS catalogues, animal models and the literature^27,28^. The PhenoScanner search of published GWAS showed that 27 of our 535 novel loci (using sentinel SNPs or proxies; r^2^ ≥ 0.8) are also strongly associated with other traits and diseases (**Fig. 5a**, **Supplementary Table 11b**). We identified *APOE* as a highly cross-related BP locus showing associations with lipid levels, cardiovascular related outcomes and Alzheimer’s disease, a finding that highlights a common link between cardiovascular risk and cognitive decline (**Fig. 5a**). Four loci overlap with anthropometric traits, including body mass index (BMI), birth weight and height. DisGeNET terms related to lipid measurements, cardiovascular outcomes and obesity overlap with BP loci **(Fig. 5b, c)**.

**Figure 5.**
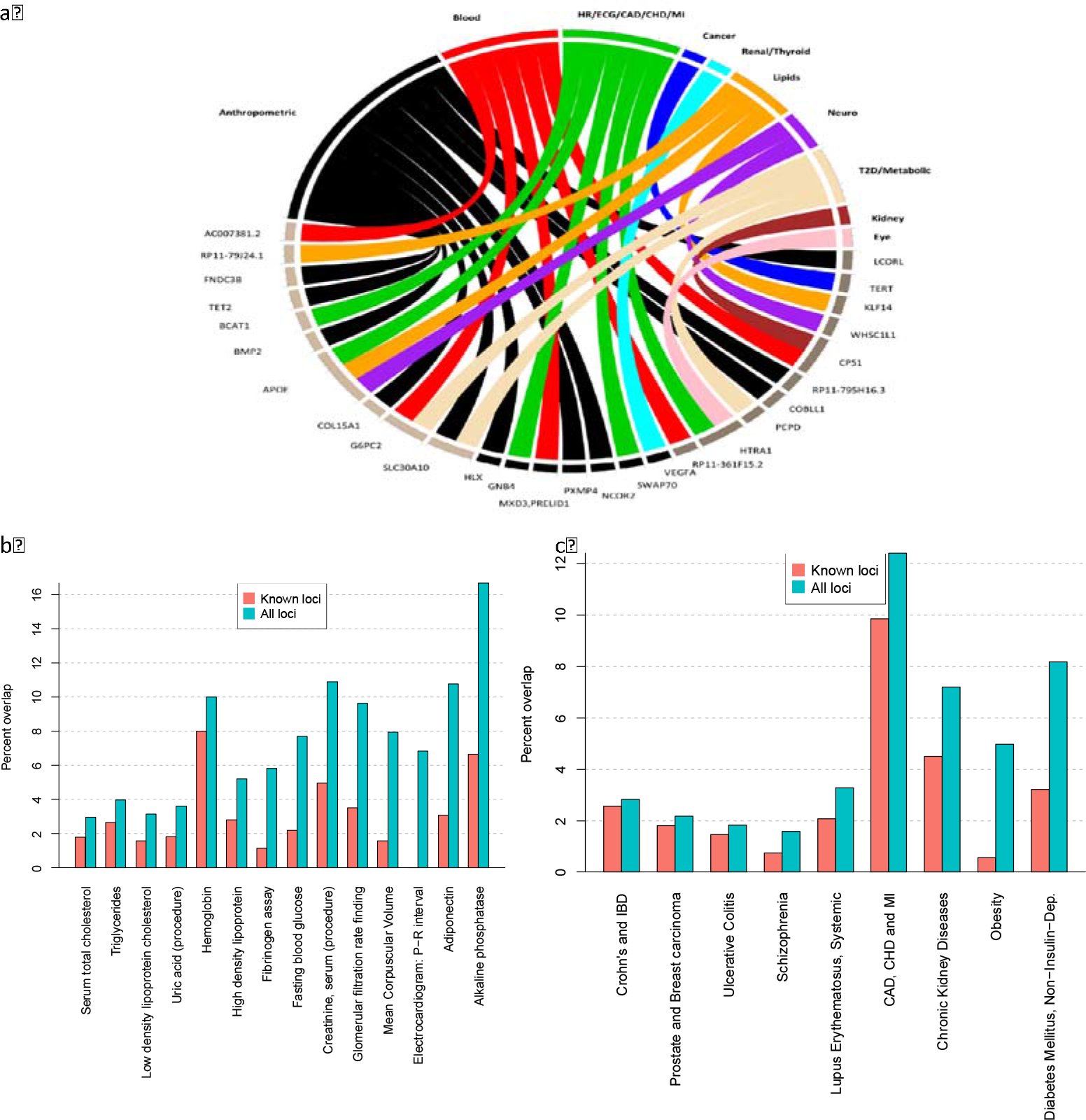
Association of blood pressure loci with other traits. Plot (a) shows results from associations with other traits which were extracted from the PhenoScanner database for the 535 novel sentinel SNPs including proxies in Linkage Disequilibrium (r^2^ ≥ 0.8) with genome-wide significant associations. Plots (b) and (c) show overlap between variants associated to (b) traits and (c) diseases in the manually-curated version of the DisGeNET database, and all variants in LD r^2^ ≥ 0.8 with SNPs from the 274 published loci (red bars), and all BP variants from all 901 loci (green bars). Diseases with an overlap of at least 5 variants with those in linkage with all markers are ordered from left to right based on how many new overlapping variants were found thanks to the validated loci. The Y-axis shows the percentage of variants associated with the diseases that is covered by the overlap. For the sake of clarity, the DisGeNET terms for blood pressure and hypertension are not displayed, whereas the following terms have been combined: CAD, CHD and MI; prostate and breast carcinoma; Crohn's and inflammatory bowel diseases. CAD: Coronary Arterial Disease; CHD: Coronary Heart Disease; MI: Myocardial Infraction

### Genetic risk of increased blood pressure, hypertension and cardiovascular disease

We created a genetic risk score (GRS) for BP levels weighted according to the effect estimates from ICBP (for published loci) and the MVP+EGCUT replication (for novel loci) across all 901 loci. The combination of these BP variants was associated with a 10.3mm Hg higher, sex-adjusted mean systolic pressure in UK Biobank for the comparison between the upper and lower quintiles of the GRS distribution (95% CI: 9.96 to 10.60 mm Hg, *P* < 1 × 10^−300^) (**Supplementary Table 12**). In addition, we observed over two-fold higher risk of hypertension (OR 2.59; 95% CI: 2.53 to 2.65; *P* < 1 × 10^−300^) in UK Biobank (**Fig. 6**). Similar results were observed comparing upper and lower deciles of the GRS distribution (12.7 mmHg difference in SBP, 95% CI: 12.2 to 13.2, *P* < 1 × 10^−300^) **(Supplementary Table 12)**. To avoid ‘winner’s curse’ we performed sensitivity analysis in the independent Airwave cohort and we observed similar results (**Supplementary Table 13**).

**Figure 6.**
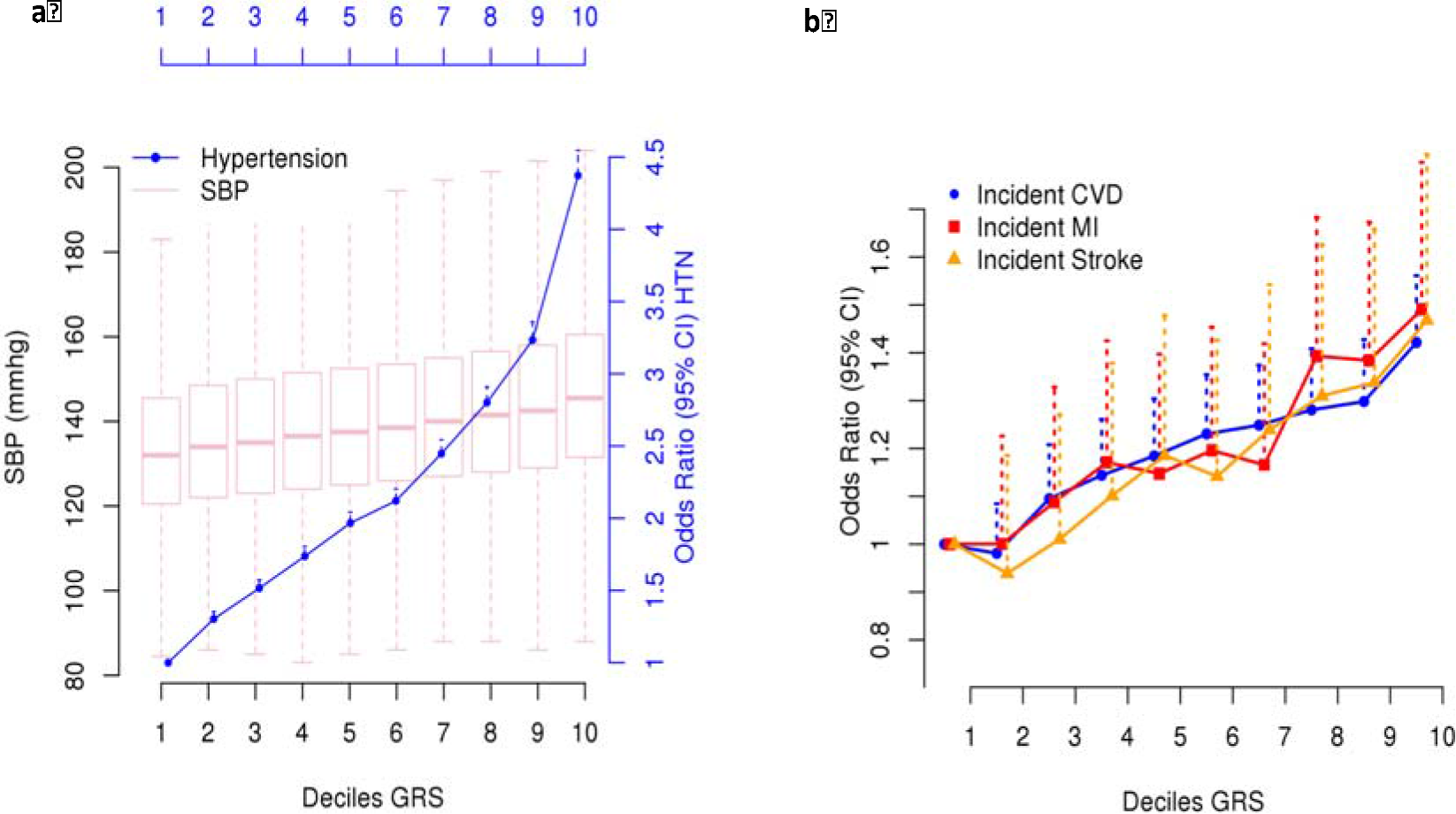
Relationship of genetic risk score (GRS) with hypertension and cardiovascular disease in UK Biobank. The GRS is based on all 901 loci, showing sex-adjusted odds ratios of (**a**) hypertension (HTN) and (**b**) incident cardiovascular disease (CVD), myocardial infarction (MI) and stroke, comparing each of the upper nine GRS deciles with the lowest decile; dotted lines represent the upper 95% confidence intervals.

Using the UK Biobank record linkage to Hospital Episode Statistics and mortality follow-up data we showed that the GRS is associated with increased risk of incident stroke, incident myocardial infarction and all incident cardiovascular outcomes, comparing the upper and lower fifths of the GRS distribution, with odds ratios of 1.37 (95% CI: 1.28 to 1.47, *P* = 1.9 × 10^−20^), 1.44 (95% CI: 1.26 to 1.64, *P* = 1.0 × 10^−7^) and 1.45 (95% CI: 1.24 to 1.68, *P* = 2.1 × 10^−6^) respectively.

### Extending analysis to other ancestries

By applying our genetic risk score to unrelated people of non-European ancestry from UK Biobank (N=6,264 Africans; N=7,881 South Asians), we show that loci associated with BP from a European population are also associated with BP in other ancestries **(Supplementary Table 14a and 14b)**. BP variants in combination were respectively associated with 6.1 mmHg (95% CI: 3.83 to 8.46) and 7.3 mmHg (95% CI: 5.14 to 9.12) higher, sex-adjusted mean systolic pressure among individuals of African and South Asian ancestries.

## DISCUSSION

Our study of over 1 million people offers a major step forward in understanding the genetic architecture of BP. The 535 novel loci we identify here have been strongly leveraged by UKB. These results more than triple the number of loci for BP traits and make major inroads into the missing heritability influencing BP level in the population^29^. These findings illustrate the power of a large-scale standardised approach to data collection, biobanking, genotyping, quality control and imputation. The novel loci open the vista of entirely new biology and highlight gene regions in systems not previously implicated in BP regulation. This is particularly timely as the global prevalence of people with SBP over 110-115 mm Hg, above which cardiovascular risk increases in a continuous graded manner, now exceeds 3.5 billion people and those within the treatment range exceed 1 billion ^30,31^.

Our functional analysis highlights the role of the vasculature and associated pathways in the genetics underpinning BP traits. For the first time, we show a role for several loci in the transforming growth factor beta (TGFβ) pathway including SMAD family genes and the *TGFfβ* gene locus itself. This pathway affects sodium handling in the kidney, ventricular remodelling and recently plasma levels of TGFβ have been correlated with hypertension (**Fig. 7**)^32,33^. The activin A receptor type 1C (*ARVC*) gene mediates the effects of the TGFβ family of signalling molecules. Another BP locus contains the Bone Morphogenetic Protein 2 (*BMP2*) gene in the TGFβ pathway, which prevents growth suppression in pulmonary arterial smooth muscle cells and is associated with pulmonary hypertension^34^. We identified another BP locus including the Kruppel-like family 14 (*KLF14*) gene of transcription factors which are induced by low levels of TGFβ receptor II gene expression. This gene has also been associated with type 2 diabetes, hypercholesterolaemia and atherosclerosis^35^.

**Figure 7:**
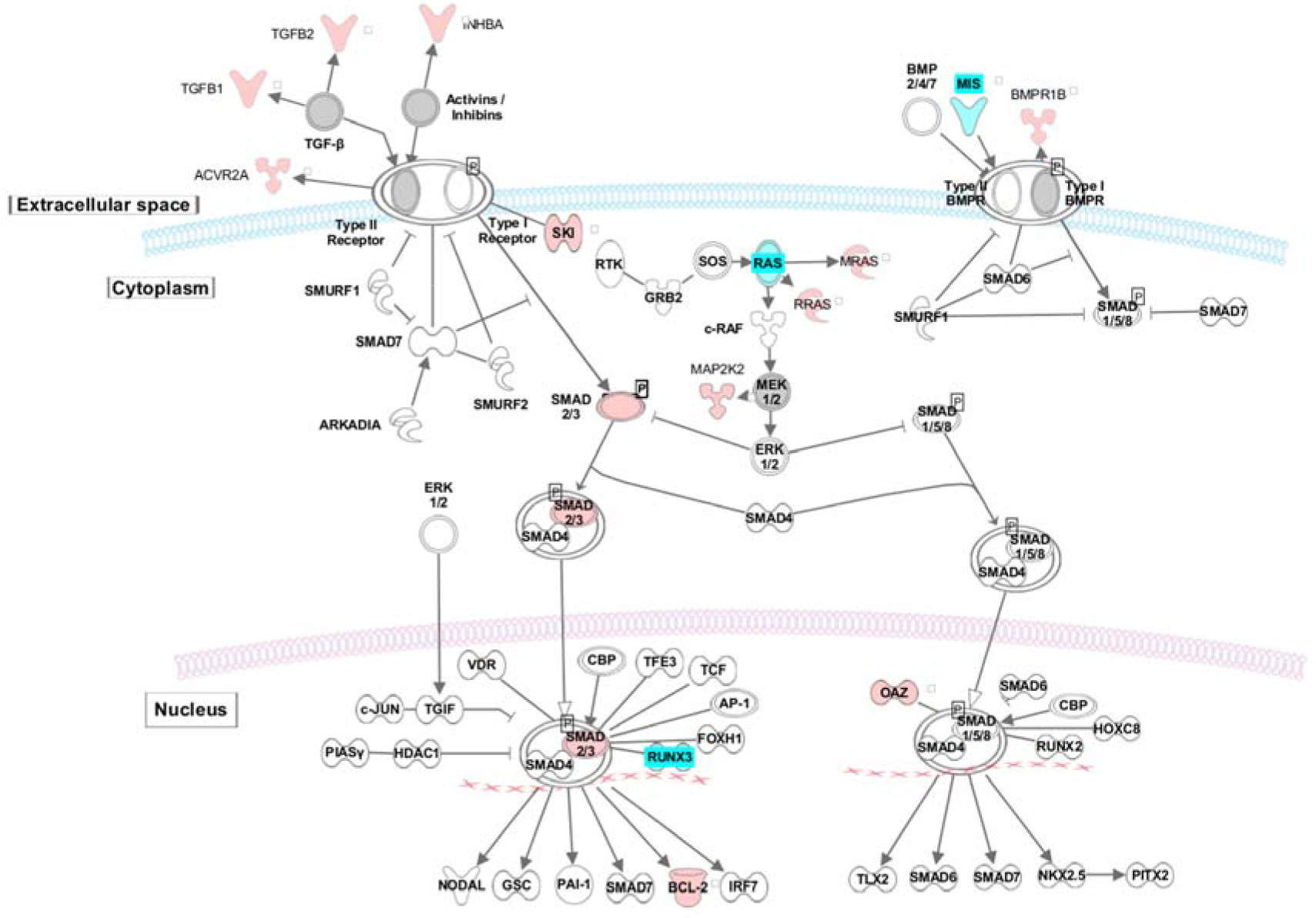
All 901 BP associations in the TGFB signalling pathway. Genes with published associations to BP are indicated in cyan. Genes with novel association to BP reported in this study are indicated in red. TGFB pathway was derived from an ingenuity canonical pathway. BP; Blood Pressure

Our analysis, for the first time, has shown a greater enrichment of BP genes in the adrenal tissue. The importance of the role of the adrenal gland in blood pressure regulation has been recognised through the observation that autonomous aldosterone production by the adrenal glands is responsible for 5-10% of all hypertension. This proportion rises further to 20% amongst people with resistant hypertension. In this analysis we add to the previously identified loci with novel loci that we can now link functionally new to aldosterone secretion^36,37^. For example the novel *CTNNB1* locus, which encodes β-catenin, the central molecule in the canonical Wnt signalling system, required for normal adrenocortical development^38,39^. Somatic adrenal mutations of this gene that prevent serine/threonine phosphorylation lead to hypertension through the generation of aldosterone-producing _adenomas_^40 41^.

Our novel loci also include genes involved in vascular remodelling including the gene for vascular endothelial growth factor A (*VEGFA*) which induces proliferation, migration of vascular endothelial cells and stimulates angiogenesis. Disruption of this gene in mice resulted in abnormal embryonic blood vessel formation. Allelic variants of this gene have been associated with microvascular complications of diabetes, atherosclerosis and the antihypertensive response to enalapril^42^. We previously reported another fibroblast growth factor (*FGF5*) gene locus in association with BP. In this study, we now identify a new BP locus encoding the FGF9 which has been linked to enhanced angiogenesis and vascular smooth muscle cell differentiation by regulating *VEGFA* expression.

Several of our novel loci contain lipid related genes which supports the observed strong associations across multiple cardio-metabolic traits. For example, the apolipoprotein E gene (*APOE*) encodes the major apoprotein of the chylomicron. Recently, APOE serum levels have been correlated with systolic BP in population-based studies and in murine knockout models’ disruption of this gene led to atherosclerosis and hypertension^43,44^. A second novel BP locus contains the low-density lipoprotein receptor-related protein 4 (*LRP4*) gene which may be a target for *APO*E and is strongly expressed in the heart in mice and humans. In addition, we identified a novel locus including the apolipoprotein L domain containing 1 gene (*APOLD1*) that is highly expressed in the endothelium of developing tissues (particularly heart) during angiogenesis.

Many of our novel BP loci encode proteins which may modulate vascular tone or signalling. For example, the locus containing urotensin-2 receptor (*UTS2R*) gene encodes a class A rhodopsin family G-protein coupled-receptor that upon activation by the neuropeptide urotensin II, produces profound vasoconstriction. One of the novel loci for SBP contains the relaxin gene which encodes a G-protein coupled receptor with roles in uterine relaxation, vasorelaxation and cardiac function and which signals by phosphatidylinositol 3-kinase (PI3K) ^45,46^. We identified the novel *RAMP2* locus which encodes an adrenomedullin receptor^47^; we previously identified the adrenomedullin (*ADM*) as a BP locus^13^. Adrenomedullin is known to exert differential effects on BP in the brain (vasopressor) and the vasculature (vasodilator). In addition, we identify the novel locus containing the *PI3K* gene encoding a signalling enzyme which inhibits vascular smooth muscle cell proliferation and neo-intimal formation^48^. A locus containing Rho guanine nucleotide exchange factor 25 (*ARHGEF25*) gene generates a factor which interacts with Rho GTPases involved in contraction of vascular smooth muscle and regulation of responses to angiotensin II^49^.

We evaluated the 901 BP loci for extant or potentially druggable targets. We note that loci encoding *MARK3*, *PDGFC*, *TRHR*, *ADORA1*, *GABRA2*, *VEGFA* and *PDE3A* are within systems that have existing drugs not currently linked to a known antihypertensive mechanism and may offer repurposing opportunities e.g. detection of *SLC5A1* as a new BP locus which is targeted by the type 2 diabetes drug Canagliflozin. This is important as between 8-12% of patients with hypertension exhibit resistance or intolerance to current therapies and repositioning of a therapy with a known safety profile may reduce development costs.

Here we demonstrate for the first time that several lifestyle exposures that are known to elevate BP may be genetically determined in part, and share that same genetic architecture with several loci that influence BP. We adjusted our BP association analyses for BMI, so whilst acknowledging the potential for collider bias between a lifestyle factor and the conditioned trait of BP, the use of adjustment should, if anything, reduce the ability to detect loci in common between two traits. Our new findings for the genetic overlap of loci associated with both BP and lifestyle exposures could allow an additional focus on specific lifestyle measures known to affect BP, offering an additional strand to a precision medicine strategy.

Our findings using a larger sample size, strengthen our previously reported genetic risk score analysis indicating that all BP elevating alleles combined could increase systolic BP by 10 mm Hg and increase risk of cardiovascular events (stroke and coronary disease)^11^. We previously suggested that by genotyping all BP elevating variants in the young that lifestyle intervention in early life might attenuate the BP rise at older ages^11^.

Our discovery of 535 novel loci from a combination of a two-stage GWAS with independent replication and one-stage GWAS design with internal replication illustrates the value of this approach to minimise the effects of stochastic variation and heterogeneity. To address the risk of false positive loci within our discovery meta-analysis we required an order of magnitude smaller *P*-value for the one-stage design in line with thresholds recommended for whole genome sequencing^22^.

The new discoveries reported here more than treble to 901 the number of loci for BP and represent a substantial advance in understanding the genetic architecture of BP in people of European descent with extension to other ancestries. Our confirmation of the impact of these variants on BP level and cardiovascular events, coupled with identification of shared risk alleles for BP and adverse lifestyle could contribute to an early life precision medicine strategy.

## Acknowledgements

H. R.W. was funded by the National Institute for Health Research (NIHR) as part of the portfolio of translational research of the NIHR Biomedical Research Centre at Barts and The London School of Medicine and Dentistry. D.M-A is supported by the Medical Research Council [grant number MR/L01632X.1]. B.M. holds an MRC eMedLab Medical Bioinformatics Career Development Fellowship, funded from award MR/L016311/1. H.G. was funded by the NIHR Imperial College Health Care NHS Trust and Imperial College London Biomedical Research Centre. C.P.C. was funded by the National Institute for Health Research (NIHR) as part of the portfolio of translational research of the NIHR Biomedical Research Center at Barts and The London School of Medicine and Dentistry. S.T. was supported by Canadian Institutes of Health Research; Université Laval (Quebec City, Canada). G.P. was supported by Canada Research Chair in Genetic and Molecular Epidemiology and CISCO Professorship in Integrated Health Biosystems. I.K. was supported by the EU PhenoMeNal project (Horizon 2020, 654241). C.P.K. is supported by grants R01DK096920 and U01DK102163 from the NIH-NIDDK, and by resources from the Memphis VA Medical Center. P.S.T. was supported by VA Award #I0BX003362. C.P.K. is an employee of the US Department of Veterans affairs. Opinions expressed in this paper are those of the authors’ and do not necessarily represent the opinion of the Department of Veterans Affairs. S.D. was supported for this work by grants from the European Research Council (ERC), the EU Joint Programme - Neurodegenerative Disease Research (JPND), the Agence Nationale de la Recherche (ANR). M.BOE is supported by NIH grant R01-DK062370. H.W. and A.G. acknowledge support of the Tripartite Immunometabolism Consortium [TrIC], Novo Nordisk Foundation (grant NNF15CC0018486). N.V. was supported by Marie Sklodowska-Curie GF grant (661395) and ICIN-NHI. C.M. is funded by the MRC AimHy (MR/M016560/1) project grant. M.A.N participation is supported by a consulting contract between Data Tecnica International and the National Institute on Aging, NIH, Bethesda, MD, USA. M.BR., M.CO., I.G., P.G., G.G, A.MO., A.R., D.V. were supported by the Italian Ministry of Health RF2010 to Paolo Gasparini, RC2008 to Paolo Gasparini. D.I.B. is supported by the Royal Netherlands Academy of Science Professor Award (PAH/6635). J.C.C. is supported by the Singapore Ministry of Health’s National Medical Research Council under its Singapore Translational Research Investigator (STaR) Award (NMRC/STaR/0028/2017). C.C., P.B.M and M.R.B were funded by the National Institutes for Health Research (NIHR) as part of the portfolio of translational research of the NIHR Biomedical Research Centre at Barts. M.R. is recipient from China Scholarship Council (No. 2011632047). C.L. was supported by the Medical Research Council UK (G1000143; MC_UU_12015/1; MC_PC_13048; MC_U106179471), Cancer Research UK (C864/A14136), EU FP6 programme (LSHM_CT_2006_037197). G.B.E is supported by the Swiss National Foundation SPUM project FN 33CM30-124087, Geneva University, and the Fondation pour Recherches Médicales, Genève. D.O.M-K. is supported by Dutch Science Organization (ZonMW-VENI Grant 916.14.023). M.M was supported by the National Institute for Health Research (NIHR) BioResource Clinical Research Facility and Biomedical Research Centre based at Guy's and St Thomas' NHS Foundation Trust and King's College London. H.W. and M.F. acknowledge the support of the Wellcome Trust core award (090532/Z/09/Z) and the BHF Centre of Research Excellence (RE/13/1/30181). A.G, H.W. acknowledge European Union Seventh Framework Programme FP7/2007-2013 under grant agreement no. HEALTH-F2-2013-601456 (CVGenes@Target) & and A.G, the Wellcome Trust Institutional strategic support fund. A.P.M. was a Wellcome Trust Senior Research Fellow in Basic Biomedical Science (WT098017). L.R. was supported by Forschungs- und Förder-Stiftung INOVA, Vaduz, Liechtenstein. P.v.d.H. was supported by ICIN-NHI and Marie Sklodowska-Curie GF (call: H2020-MSCA-IF-2014, Project ID: 661395). . N.J.W. was supported by the Medical Research Council UK (G1000143; MC_UU_12015/1; MC_PC_13048; MC_U106179471), Cancer Research UK (C864/A14136), EU FP6 programme (LSHM_CT_2006_037197). E.Z. was supported by the Wellcome Trust (WT098051). J.N.H. and A.G. were supported by VA grant 1I01CX000982. T.L.E. and D.R.V.E. were supported by grant R21HL121429 from NIH/NHLBI. A.M.H. was supported by VA Award #I01BX003360. C.J.O. was supported by the VA Boston Healthcare, Section of Cardiology and Department of Medicine, Brigham and Women’s Hospital, Harvard Medical School. L.V.W. holds a GlaxoSmithKline/British Lung Foundation Chair in Respiratory Research. P.E. acknowledges support from the NIHR Biomedical Research Centre at Imperial College Healthcare NHS Trust and Imperial College London, the NIHR Health Protection Research Unit in Health Impact of Environmental Hazards (HPRU-2012-10141), and the Medical Research Council (MRC) and Public Health England (PHE) Centre for Environment and Health (MR/L01341X/1). P.E. is a UK Dementia Research Institute (DRI) professor, UK DRI at Imperial College London, funded by the MRC, Alzheimer’s Society and Alzheimer’s Research UK. M.J.C. was funded by the National Institute for Health Research (NIHR) as part of the portfolio of translational research of the NIHR Biomedical Research Center at Barts and The London School of Medicine and Dentistry. M.J.C. is a National Institute for Health Research (NIHR) senior investigator.

This research was supported by the British Heart Foundation (grant SP/13/2/30111). Project title: Large-Scale Comprehensive Genotyping of UK Biobank for Cardiometabolic Traits and Diseases: UK CardioMetabolic Consortium (UKCMC). This research has been conducted using the UK Biobank Resource under application number 236.

Computing: This work was enabled using the computing resources of the i) UK MEDical BIOinformatics partnership - aggregation, integration, visualisation and analysis of large, complex data (UK MED-BIO) which is supported by the Medical Research Council [grant number MR/L01632X/1] and ii) the MRC eMedLab Medical Bioinformatics Infrastructure, supported by the Medical Research Council [grant number MR/L016311/1].

## Conflicts/Disclosures

M.A.N. consults for Illumina Inc, the Michael J. Fox Foundation and University of California Healthcare among others.

M.J.C. is Chief Scientist for Genomics England, a UK Government company.

The views expressed in this manuscript are those of the authors and do not necessarily represent the views of the National Heart, Lung, and Blood Institute; the National Institutes of Health; or the U. S. Department of Health and Human Services. This publication does not represent the views of the Department of Veterans Affairs or the United States Government.

## Author contributions

***Central analysis:*** E.E., H.R.W., D.M-A., B.M., R.P., H.G., G.N., N.D., C.P.C., I.K., F.N., M E., K.W., E.T. L.V.W.

***Writing of the manuscript:*** E.E., H.R.W., D.M-A., B.M., R.P., H.G., IT., M.R.B., L.V.W., P.E., M.J.C. (with group leads EE, H.R.W, L.V.W., P.E., M.J.C.)

***ICBP-Discovery contributor:*** (3C-Dijon) S.D., M.S., P.AM., G.C., C.T.; (AGES-Reykjavik) V.GU., L.J.L., A.V.S., T.B.H.; (ARIC) D.E.A., E.B., A.CH. A.C.M., P.N.; (ASCOT) N.R.P., D.C.S., A S., S.THO., P.B.M., P.S., M.J.C., H.R.W.; (ASPS) E.H., Y.S., R.S., H.S.; (B58C) D.P.S., BHSA.J., N.SHR.; (BioMe (formerly IPM)) E.P.B., Y.LU., R.J.F.L.; (BRIGHT) J.C., M.F., M.J.B., P.B.M., M.J.C., H.R.W.; (CHS) J.C.B., K.R., K.D.T., BMP.; (Cilento study) M.C., T.NU., D R., R.SO.; (COLAUS) M B., Z.K., P.V.; (CROATIA_Korcula) J.MART., A.F.W.; (CROATIA_SPLIT) I.KO., O.P., T.Z.; (CROATIA_Vis) J.E.H., I.R., V.V.; (EPIC) K-T.K., R.J.F.L., N.J.W.; (EPIC-CVD) W-Y.L., P.SU., A.S.B., J.DA., J.M.M.H.; (EPIC-Norfolk, Fenland-OMICS, Fenland-GWAS) J-H.Z.; (EPIC-Norfolk, Fenland-OMICS, Fenland-GWAS, InterAct-GWAS) J.L., C.L., R.A.S., N.J.W.; (ERF) N.A., B.A.O., C.M.v.D.; (Fenland-Exome, EPIC-Norfolk-Exome) S.M.W., FHSS-J.H., D.L.; (FINRISK (COROGENE_CTRL)) P.J., K.K., M.P., A-P.S.; (FINRISK_PREDICT_CVD) A.S.H., A.P., S R., V S.; (FUSION) A.U.J, M.BOE., F.C., J.T., (GAPP) S T., G.P., D.CO., L.R.; (Generation Scotland (GS:SFHS)) T.B., C.H., A.C., S.P.; (GoDARTs) N.S., A.S.F.D., A.D.M., C.N.A.P.; (GRAPHIC) P.S.B., C.P.N., N.J.SA., M.D.T.; (H2000_CTRL) A.JU., P.K., S.KO., T.N.; (HABC) Y.L., M.A.N., T.B.H.; (HCS) E.G.H., C O., R.J.SC.; (HTO) K.L.A., H.J.C., B.D.K., C.MA.; (ICBP-SC) G.A., T.F., M-R.J., A D J., M.LA., C.N.; (INGI-CARL) I.G., G.G., A MO., A.R.; (INGI-FVG) M.BR., M.CO., P.G., D.V.; (INGI-VB) C. M.B., N.J.S., D.T., M.T.; (JUPITER) F.G., L.M.R., P.M.R., D.I.C.; (KORA S3) C.G., M L., E.O., S.S.; (KORA S4) A PE., J.S.R.; (LBC1921) S.E.H., D.C.M.L., A.PA., J.M.S.; (LBC1936) G.D., I.J.D., A.J.G., L.M.L.; (Lifelines) N.V., M.H.d.B., M A S., P.v.d.H.; (LOLIPOP) J.C.C., J.S.K., B.L., W.Z.; (MDC) P.A., O.M.; (MESA) X.G., W.P., J.I.R., J.Y.; (METSIM) A.U.J., M.LAA.; (MICROS) F.M.D., A.A.H., P.P.P.; (MIGEN) R.E., S.K., J.M., D. SI.; NEO) R.L., R.d.M., R.N., D.O.M-K.; (NESDA) Y.M., I.M.N., B.W.J.H.P., H.SN.; (NSPHS) S.E., U.G., Å.JO.; (NTR) D.I.B., E.J.d.G., J-J.H., G.W.; (ORCADES) H.C., P.K.J., S.H.W., J.F.W.; (PIVUS) L.LI., C.M.L., J.S., A.M.; (Prevend) N.V., P.v.d.H.; (PROCARDIS) M.F., A.G., H.W.; (PROSPER) J.DE., J.W.J., D.J.S., S.TR.; (RS) O.H.F., A.HO., A.U., G.C.V.; (SardiNIA) J.D., Y.Q., F.CU., E.G.L.; (SHIP) M.D., R.R., A T., U.V.; (STR) M.FR., A.H., R.J.S., E.I.; (TRAILS) C.A.H., A.J.O., H R., P.J.v.d.M.; (TwinsUK) M.M., C M., T.D.S.; (UKHLS) B.P., E.Z.; (ULSAM) V.G., A.P.M., A.M., E.I.; (WGHS) F.G., L.M.R., P.M.R., D.I.C.; (YFS) M.K., T.L., L-P.L., O.T.R.

***Replication study contributor:*** (MVP) J.N.H., A.G., D.R.V.E., Y.V.S., K.C., J.M.G., P.W.F.W., P.S.T., C.P.K., A.M.H., C.J.O., T.L.E.; (EGCUT) T.E., R.M., L.M. A.ME.

***Airwave Health Monitoring Study:*** E.E, H.G, A-C.V., R.P., I.K., I T., P.E.

***All authors critically reviewed and approved the final version of the manuscript***

## METHODS

### UK Biobank (UKB) data

We performed a Genome Wide Association Study (GWAS) analysis in 458,577 UKB participants^14^ (**Supplementary Methods**). These consist of ~408,951 individuals from UKB genotyped at 807,411 variants with a custom Affymetrix UK Biobank Axiom Array chip and 49,626 individuals genotyped at 825,927 variants with a custom Affymetrix UK BiLEVE Axiom Array chip from the UK BiLEVE study^50^, which is a subset of UKB. SNPs were imputed centrally by UKB using a reference panel that merged the UK10K and 1000 Genomes Phase 3 panel as well as the Haplotype Reference Consortium (HRC) panel^51^. For current analysis only SNPs imputed from the HRC panel were considered.

#### UKB phenotypic data

Following Quality Control (QC) **(Supplementary Methods)**, we restricted our data to a subset of post-QC individuals of European ancestry combining information from selfreported and genetic data **(Supplementary Methods)** resulting in a maximum of N=458,577 individuals **(Fig. 1**, **Supplementary Fig. 9)**.

Three BP traits were analysed: systolic (SBP), diastolic (DBP) and pulse pressure (PP) (difference between SBP and DBP). We calculated the mean SBP and DBP values from two automated (N=418,755) or two manual (N=25,888) BP measurements. For individuals with one manual and one automated BP measurement (N=13,521), we used the mean of these two values. For individuals with only one available BP measurement (N=413), we used this single value. After calculating BP values, we adjusted for medication use by adding 15 and 10 mmHg to SBP and DBP, respectively, for individuals reported to be taking BP-lowering medication (94,289 individuals)^52^. Descriptive summary statistics are shown in **Supplementary Table 1a.**

#### UKB Analysis models

For the UKB GWAS we performed linear mixed model (LMM) association testing under an additive genetic model of the three (untransformed) continuous, medication-adjusted BP traits (SBP, DBP, PP) for all measured and imputed genetic variants in dosage format using the BOLT-LMM (v2.3) software^16^. We used genotyped SNPs filtered for MAF > 5%; HWE *P* > 1×10^−6^; missingness < 0.015, to estimate the parameters of the linear mixed model, for the initial modelling step only. Within the association analysis, we adjust for the following covariates: sex, age, age^2^, BMI and a binary indicator variable for UKB vs UK BiLEVE to account for the different genotyping chips and different study ascertainment. The association analysis performed by BOLT-LMM (v2.3) corrects for population structure and cryptic relatedness in very large datasets^16^. The genome-wide association analysis of all HRC-imputed SNPs was restricted to variants with MAF ≥ 1% and INFO > 0.1.

#### Genomic inflation and confounding

We applied the univariate LD score regression method (LDSR)^17^ to test for genomic inflation (expected for polygenic traits like BP, with large sample sizes, and especially also from analyses of such dense genetic data with so many SNPs in high-LD)^53^. LDSR intercepts (and standard errors) were 1.217 (0.018), 1.219 (0.020) and 1.185 (0.017) for DBP, SBP and PP respectively, and were used to adjust the UKB GWAS results for genomic inflation, prior to the meta-analysis.

### International Consortium for Blood Pressure (ICBP) GWAS

ICBP GWAS is an international consortium to investigate BP genetics^7,12,54^. We combined previously reported post-quality control (QC) GWAS data from 54 studies (N=150,134)^12^, with newly available GWAS data from a further 23 independent studies (N=148,890) using a fixed effects inverse variance weighted meta-analysis. The 23 studies providing new data were: ASCOT-SC, ASCOT-UK, BRIGHT, Dijon 3C, EPIC-CVD, GAPP, HCS, GS:SFHS, Lifelines, JUPITER, PREVEND, TWINSUK, GWAS-Fenland, InterAct-GWAS, OMICS-EPIC, OMICS-Fenland, UKHLS, GoDARTS-Illumina and GoDarts-Affymetrix, NEO, MDC, SardiNIA, METSIM.

All study participants were of European descent and were imputed to either the 1000 Genomes Project Phase 1 integrated release v.3 [March 2012] all ancestry reference panel^55^ or the Haplotype Reference Consortium (HRC) panel^15^. The final enlarged ICBP GWAS dataset included 77 cohorts (N=299,024 individuals).

Definition of phenotype data and GWAS analyses of SBP, DBP and PP were as per previous ICBP protocol for 54 studies^12^, extended to the additional 23 studies for which new data were available. Full study names, cohort information and general study methods are included in **Supplementary Table 1b** and in Supplementary **Tables 15a-c**. Genomic control was applied at study-level. The LDSR intercepts (standard error) for the ICBP GWAS meta-analysis were 1. 089 (0.012), 1.086 (0.012) and 1.066 (0.011) for SBP, DBP and PP, respectively.

### Meta-analyses of discovery datasets

We performed a fixed-effects inverse variance weighted meta-analysis using METAL^19,56^ to obtain summary results from the combined UKB and ICBP GWAS, for up to N=757,601 participants and ~7.1 M SNPs with MAF ≥ 1% present in both the HRC-imputed UKB data and ICBP meta-analysis for all three traits. The LDSR intercepts (standard error), in the discovery meta-analysis of UKB and ICBP were 1.156 (0.020), 1.160 (0.021) and 1.113 (0.018) for SBP, DBPP and PP respectively. The LDSR intercept (standard error), after the exclusion of all published BP variants (see below) in the discovery meta-analysis of UKB and ICBP was 1.090 (0.018), 1.097 (0.017) and 1.064 (0.015) for SBP, DBP and PP respectively; hence showing very little inflation in the discovery GWAS after the exclusion of published loci **(Supplementary Fig. 10).** No further correction was applied to the discovery meta-analysis of UKB and ICBP GWAS.

### Previously reported variants

We compiled from the peer-reviewed literature all 357 SNPs previously reported to be associated with BP at the time that our analysis was completed, that have been identified and validated as the sentinel SNP in primary analyses from previous BP genetic association studies. These 357 published SNPs correspond to 274 distinct loci, according to locus definition of: (i) SNPs within ±500kb distance of each other; (ii) SNPs in Linkage Disequilibrium (LD), using a threshold of r^2^ ≥ 0.1, calculated with PLINK (v2.0). We then augment this list to all SNPs present within our data, which are contained within these 274 published BP loci, i.e. all SNPs which are located ±500kb from each of the 357 published SNPs and/or in LD with any of the 357 previously validated SNPs (r^2^ ≥ 0.1). The LDSR intercept (standard error), after the exclusion of all published BP variants in the discovery meta-analysis of UKB and ICBP was 1.090 (0.018), 1.097 (0.017) and 1.064 (0.0146) for SBP, DBP and PP respectively; hence showing very little inflation in the discovery GWAS after the exclusion of published loci **(Supplementary Fig. 10).**

### Identification of novel signals: two-stage and one-stage study designs

To identify novel signals of association with BP, two complementary study designs (which we term here “two-stage design” and “one-stage design”) were implemented in order to maximise the available data and minimise reporting of false positive associations.

### Two-stage design: Overview

All of the following criteria must be satisfied for a signal to be reported as a novel signal of association with BP using our two-stage design:

i. the sentinel SNP shows significance (*P* < 1 × 10^−6^) in the discovery meta-analysis of UKB and ICBP, with concordant direction of effect between UKB and ICBP;
ii. the sentinel SNP is genome-wide significant (***P* < 5 × 10**^**−8**^) in the combined meta analysis of discovery and replication (MVP and EGCUT) (replication, described below);
iii. the sentinel SNP shows support (*P* < 0.01) in the replication meta-analysis of MVP and EGCUT alone **(Supplementary Methods)**;
iv. the sentinel SNP has concordant direction of effect between the discovery and the replication meta-analyses;
v. the sentinel SNP must not be located within any of the 274 previously reported loci described above.

The primary replicated trait was then defined as the replicated BP trait with the most significant association from the combined meta-analysis of discovery and replication (in the case of many SNPs replicating for more than one BP trait).

### Two-stage design: selection of variants from the discovery meta-analysis

We considered for follow-up SNPs in loci non-overlapping with previously reported loci according to both an LD threshold of r^2^ ≥ 0.1 and a 1Mb interval region, as calculated with PLINK^57^. We obtained a list of such SNPs with *P* < 1×10^−6^ for any of the three BP traits, which also had concordant direction of effect between UKB vs ICBP. By ranking the SNPs by significance in order of minimum P-value across all BP traits, we performed an iterative algorithm to determine the number of novel signals **(Supplementary Methods)**, and identify the sentinel SNP (most significant) per locus.

### Two-stage design: replication analysis

We used two independent external data sets for replication **(Supplementary Methods)**. We considered SNPs with MAF ≥ 1% for an independent replication in MVP (max N = 220,520)^20^ and in EGCUT Biobank (N=28,742)^21^. This provides a total of N = 249,262 independent samples of European descent available for replication. Additional information on the analyses of the two replication datasets is provided in **Supplementary Methods** and in **Supplementary Table 1c**.

The two datasets were then combined using fixed effects inverse variance weighted metaanalysis and summary results for all traits were obtained for the replication meta-analysis dataset.

### Two-stage design: combined meta-analysis of discovery and replication meta-analyses

The meta-analyses were performed within METAL software^56^ using fixed effects inverse variance weighted meta-analysis **(Supplementary Methods)**. The combined meta-analysis of both the discovery data (N = 757,601) and replication meta-analysis (max N = 249,262) provided a maximum sample size of N = 1,006,863.

### One-stage design: Overview

All of the following criteria must be satisfied for a signal to be reported as a novel signal of association with BP using our one-stage criteria:

i. the sentinel SNP has **P < 5 × 10**^**−9**^ in the discovery (UKB+ICBP) meta-analysis;
ii. the sentinel SNP shows support (*P* < 0.01) in the UKB GWAS alone;
iii. the sentinel SNP shows support (*P* < 0.01) in the ICBP GWAS alone;
iv. the sentinel SNP has concordant direction of effect between UKB and ICBP datasets;
v. The sentinel SNP must not be located within any of the 274 previously reported loci described above or the recently reported non-replicated loci from Hoffman et al^10^.

We selected the one-stage *P*-value threshold to be more stringent than a genome-wide significance *P*-value, in order to ensure robust findings and to minimize false positives. The threshold of *P* < 5 × 10^−9^ has been proposed as a more conservative statistical significance threshold, e.g. for whole-genome sequencing-based studies^22^.

Selection of variants from the meta-analysis of UKB and ICBP was performed as described above for the two-stage design. Signals of association which met both the two-stage and one-stage study design criteria are reported using the two-stage design results only.

### Metabolomics lookups

We carry out metabolomics analysis using three sets of data. First, we use ^1^H NMR lipidomics data on plasma from a subset of 2,022 participants of the Airwave Health Monitoring Study (**Supplementary Methods**). For each novel BP-associated SNP we ran association tests with the lipidomics data using linear regression analyses, adjusted for age and sex. We computed significance thresholds using a Bonferroni correct *P*-value (4.7 × 10^−4^). We also examined associations between each novel SNP and a subset of 1,941 participants of the Airwave Health Monitoring Study with data from Metabolon platform. Finally, we also test each novel SNP (and proxy SNPs with an r^2^ ≥ 0.8) against published genome-wide vs metabolome-wide associations in plasma and urine using publicly available data from the “PhenoScanner Server”^26^ to identify metabolites that have been associated with variants of interest at *P* < 5 × 10^−8^.

### Functional analyses: Variants

We used an integrative bioinformatics approach to collate functional annotation at both the variant level (for each sentinel SNP within all BP loci) and the gene level (using SNPs in LD r^2^ ≥ 0.8 with the sentinel SNPs). At the variant level, we use Variant Effect Predictor (VEP) to obtain comprehensive characterization of variants, including consequence (e.g. downstream or non-coding transcript exon), information on nearest genomic features and, where applicable, amino acid substitution functional impact, based on SIFT and PolyPhen. The biobaRt R package is used to further annotate the nearest genes.

We evaluate all SNPs in LD (r^2^ ≥ 0.8) with our novel sentinel SNPs for evidence of mediation of expression quantitative trait loci (eQTL) in all 44 tissues using the Genotype-Tissue Expression (GTEx) database, to highlight specific tissue types which show eQTLs for a larger than expected proportion of novel loci. We further seek to identify novel loci with the strongest evidence of eQTL associations in arterial tissue, in particular.

We annotated nearest genes, eGenes (genes whose expression is affected by eQTLs) and Hi-C interactors with HUVEC, HVSMC and HAEC expression from the Fantom5 project. Genes that had higher then median expression levels in the given cell types were indicated as expressed.

To identify SNPs in the novel loci that have a non-coding functional effect (influence binding of transcription factors or RNA polymerase, or influence DNase hypersensitivity sites or histone modifications), we used DeepSEA, a deep learning algorithm, that learnt the binding and modification patterns of ~900 cell/factor combinations^58^. A change >0.1 in the binding score predicted by DeepSEA for the reference and alternative alleles respectively has been shown to have high true positive rate ~80-95% and low false positive rate ~5-10% therefore we used this cut-off to find alleles with non-coding functional effect.

We identify potential target genes of regulatory SNPs using long-range chromatin interaction (Hi-C) data from HUVECs^23^, aorta, adrenal glands, neural progenitor and mesenchymal stem cell, which are tissues and cell types that are considered relevant for regulating BP^24^. Hi-C data is corrected for genomic biases and distance using the Hi-C Pro and Fit-Hi-C pipelines according to Schmitt et al. (40kb resolution – correction applied to interactions with 50kb-5Mb span)^24^. We find the most significant promoter interactions for all potential regulatory SNPs (RegulomeDB score ≤ 5) in LD (r^2^ ≥ 0.8) with our novel sentinel SNPs and published SNPs, and choose the interactors with the SNPs of highest regulatory potential to annotate the loci.

We then perform overall enrichment testing across all loci. Firstly, we use DEPICT^59^ (Data-driven Expression Prioritized Integration for Complex Traits) to identify highly expressed tissues and cells within the BP loci. DEPICT uses a large number of microarrays (~78k) to identify cells and tissues where the genes are highly expressed and uses pre-computed GWAS phenotypes to adjust for co-founding sources. Secondly, we use DEPICT to test for enrichment in gene sets associated with biological annotations (manually curated and molecular pathways, phenotype data from mouse KO studies). Using the co-expression data DEPICT calculates a probability for each gene to belong to a given gene set and uses this to weight the enrichment of the genes present in the tested loci. DEPICT provides a *P*-value of enrichment and false discovery rates adjusted *P*-values for each tissue/cells or gene set tested. We report significant enrichments with a false discovery rate <0.01. The variants tested were the 357 published BP associated SNPs at the time of analysis and a set including all (published and novel) (for novel: combined *P* < 1 × 10^−12^) variants.

Furthermore, to investigate cell type specific enrichment within DNase I sites, we used FORGE, which tests for enrichment of SNPs within DNase I sites in 123 cell types from the Epigenomics Roadmap Project and ENCODE^60^ **(Supplementary Methods)**. Two analyses were compared (i) using published sentinel SNPs only; (ii) using sentinel SNPs at all 901 loci in order to evaluate the overall tissue specific enrichment of BP associated variants.

### Functional Analyses: Genes

At the gene level, we use Ingenuity Pathway Analysis (IPA) software (IPA®, QIAGEN Redwood City) to review genes with prior links to BP, based on annotation with the “Disorder of Blood Pressure”, “Endothelial Development” and “Vascular Disease” Medline Subject Heading (MESH) terms. We used the Mouse Genome Informatics (MGI) tool to identify BP and cardiovascular relevant mouse knockout phenotypes for all genes linked to BP in our study. We also used IPA to identify genes that interact with known targets of anti-hypertensive drugs. Genes were also evaluated for evidence of small molecule druggability or known drugs based on queries of the Drug Gene Interaction database.

### Effects on other traits and diseases

We query SNPs against PhenoScanner^26^ to investigate cross-trait effects, extracting all association results with genome-wide significance at *P* < 5 × 10^−8^ for all SNPs in high LD (r^2^ ≥ 0.8) with the 535 sentinel novel SNPs, to highlight the loci with strongest evidence of association with other traits. We further evaluated these effects using DisGeNET, a resource that integrates data from expert curated repositories, GWAS catalogues, animal models and the literature^27,28^ Specifically, at the SNP level, overlaps with DisGeNET terms were computed, with roughly the same number of markers in the published and novel BP loci. Thus, given the expected saturation of the overlaps, a more than double increase indicates that strong associations are more frequent in the novel BP loci. At the gene level, overrepresentation enrichment analysis (ORA) with WebGestalt^61^ on the nearest genes to all BP loci was carried out. Moreover, we tested sentinel SNPs at all published and novel (n=901) loci for association with lifestyle related data including food, water and alcohol intake, anthropomorphic traits and urinary sodium, potassium and creatinine excretion using the recently developed Stanford Global Biobank Engine and the Gene ATLAS^62^. Both are search engines for GWAS findings for multiple phenotypes in UK Biobank. We used a Bonferroni corrected significance threshold of *P* < 1×10^−6^ to deem significance.

### Genetic risk scores and percentage of variance explained

We calculated a genetic risk score (GRS) to provide an estimate of the combined effect of the BP raising variants on BP and risk of hypertension, and applied this to the UK Biobank data. We first create two trait-specific weighted GRSs (i.e. SBP, DBP), for all pairwise-independent, LD-filtered (r^2^ < 0.1) previously reported variants and 535 novel sentinel variants combined. For the previously reported variants, we weight BP increasing alleles by the trait-specific beta coefficients from the ICBP meta-analysis GWAS that is part of the discovery stage. For the novel variants, beta coefficients of the replication meta-analysis for each BP trait are used as independent, unbiased weights. We then derive a single BP GRS as the average of the GRS for SBP and DBP, and standardize it to have mean zero and standard deviation of one. We assess the association of the continuous GRS variable on BP by simple linear regression, and use logistic regression to examine the association of the GRS with risk of hypertension, with and without adjustment for sex. We then use linear and logistic regression to compare BP levels and risk of hypertension, respectively, for individuals in the top vs bottom quintiles of the GRS distribution. Similar analyses were performed for the top vs bottom deciles of the GRS distribution. All analyses were restricted to unrelated individuals of European ancestry (n= 292,347) from UKB. As a sensitivity analysis to avoid winner's curse bias we have applied similar analyses in Airwave, an independent cohort of N=14,004 unrelated participants of European descent **(Supplementary Methods)**.

We also assessed the association of the GRS with cardiovascular disease in unrelated participants in UKB data, based on self-reported medical history, and linkage to hospitalization and mortality data. We use logistic regression with binary outcome variables for composite incident cardiovascular disease **(Supplementary Methods)**, incident myocardial infarction and incident stroke (using the algorithmic UKB definitions) and GRS as explanatory variable (with and without sex adjustment).

We also calculated the association of this GRS with BP in unrelated African (N=6,264) and South Asian (N=7,881) samples from the UKB using the approach described above, to see whether BP-associated SNPs identified from GWAS predominantly in Europeans area also associated with BP in populations of non-European ancestry.

To calculate the percent of variance in BP explained by genetic variants in Airwave (N=14,004), we generated the residuals from a regression of each trait against age, age^2^, sex and BMI. We then fit a second linear model for the trait residuals with i) the published variants and ii) all the variants in the GRS plus the top 10 principal components, and estimate the percentage variance of the dependent (BP) variable explained by the GRS for the published and for all loci.

### Data availability statement

The genetic and phenotypic UK Biobank data are available upon application to the UK Biobank (https://www.ukbiobank.ac.uk). All replication data generated during this study are included in the published article. For example, association results of look-up variants from our replication analyses and the subsequent combined meta-analyses are contained within all Supplementary Tables provided.

### Ethics Statement

The UKB study has approval from the North West Multi-Centre Research Ethics Committee. Any participants from UKB who withdrew consent have been removed from our analysis. Each cohort within ICBP meta-analysis as well as our independent replication cohorts of MVP and EGCUT had ethical approval locally. More information for the participating cohorts is available at the Supplementary Methods.

## References

1. Forouzanfar, M.H. et al. Global Burden of Hypertension and Systolic Blood Pressure of at Least 110 to 115 mm Hg, 1990-2015. JAMA 317, 165–182 (2017).

2. Munoz, M. et al. Evaluating the contribution of genetics and familial shared environment to common disease using the UK Biobank. Nat Genet 48, 980–3 (2016).

3. Poulter, N.R., Prabhakaran, D. & Caulfield, M. Hypertension. Lancet 386, 801–12 (2015).

4. Mongeau, J.G., Biron, P. & Sing, C.F. The influence of genetics and household environment upon the variability of normal blood pressure: the Montreal Adoption Survey. Clin Exp Hypertens A 8, 653–60 (1986).

5. Feinleib, M. et al. The NHLBI twin study of cardiovascular disease risk factors: methodology and summary of results. Am J Epidemiol 106, 284–5 (1977).

6. Cabrera, C.P. et al. Exploring hypertension genome-wide association studies findings and impact on pathophysiology, pathways, and pharmacogenetics. Wiley Interdiscip Rev Syst Biol Med 7, 73–90 (2015).

7. Ehret, G.B. et al. The genetics of blood pressure regulation and its target organs from association studies in 342,415 individuals. Nat Genet 48, 1171–1184 (2016).

8. Surendran, P. et al. Trans-ancestry meta-analyses identify rare and common variants associated with blood pressure and hypertension. Nat Genet 48, 1151–1161 (2016).

9. Liu, C. et al. Meta-analysis identifies common and rare variants influencing blood pressure and overlapping with metabolic trait loci. Nat Genet 48, 1162–70 (2016).

10. Hoffmann, T.J. et al. Genome-wide association analyses using electronic health records identify new loci influencing blood pressure variation. Nat Genet 49, 54–64 (2017).

11. Warren, H.R. et al. Genome-wide association analysis identifies novel blood pressure loci and offers biological insights into cardiovascular risk. Nat Genet 49, 403–415 (2017).

12. Wain, L.V. et al. Novel Blood Pressure Locus and Gene Discovery Using Genome-Wide Association Study and Expression Data Sets From Blood and the Kidney. Hypertension (2017).

13. International Consortium for Blood Pressure Genome-Wide Association Studies et al. Genetic variants in novel pathways influence blood pressure and cardiovascular disease risk. Nature 478, 103–9 (2011).

14. Sudlow, C. et al. UK biobank: an open access resource for identifying the causes of a wide range of complex diseases of middle and old age. PLoS Med 12, e1001779 (2015).

15. McCarthy, S. et al. A reference panel of 64,976 haplotypes for genotype imputation. Nat Genet 48, 1279–83 (2016).

16. Loh, P.R. et al. Efficient Bayesian mixed-model analysis increases association power in large cohorts. Nat Genet 47, 284–90 (2015).

17. Bulik-Sullivan, B.K. et al. LD Score regression distinguishes confounding from polygenicity in genome-wide association studies. Nat Genet 47, 291–5 (2015).

18. Ioannidis, J.P., Patsopoulos, N.A. & Evangelou, E. Heterogeneity in meta-analyses of genome-wide association investigations. PLoS One 2, e841 (2007).

19. Evangelou, E. & Ioannidis, J.P. Meta-analysis methods for genome-wide association studies and beyond. Nat Rev Genet 14, 379–89 (2013).

20. Gaziano, J.M. et al. Million Veteran Program: A mega-biobank to study genetic influences on health and disease. J Clin Epidemiol 70, 214–23 (2016).

21. Leitsalu, L. et al. Cohort Profile: Estonian Biobank of the Estonian Genome Center, University of Tartu. Int J Epidemiol 44, 1137–47 (2015).

22. Pulit, S.L., de With, S.A. & de Bakker, P.I. Resetting the bar: Statistical significance in whole-genome sequencing-based association studies of global populations. Genet Epidemiol 41, 145–151 (2017).

23. Rao, S.S. et al. A 3D map of the human genome at kilobase resolution reveals principles of chromatin looping. Cell 159, 1665–80 (2014).

24. Schmitt, A.D. et al. A Compendium of Chromatin Contact Maps Reveals Spatially Active Regions in the Human Genome. Cell Rep 17, 2042–2059 (2016).

25. Elliott, P. et al. The Airwave Health Monitoring Study of police officers and staff in Great Britain: rationale, design and methods. Environ Res 134, 280–5 (2014).

26. Staley, J.R. et al. PhenoScanner: a database of human genotype-phenotype associations. Bioinformatics 32, 3207–3209 (2016).

27. Pinero, J. et al. DisGeNET: a comprehensive platform integrating information on human disease-associated genes and variants. Nucleic Acids Res 45, D833–D839 (2017).

28. Pinero, J. et al. DisGeNET: a discovery platform for the dynamical exploration of human diseases and their genes. Database (Oxford) 2015, bav028 (2015).

29. Ehret, G.B. & Caulfield, M.J. Genes for blood pressure: an opportunity to understand hypertension. Eur Heart J34, 951–61 (2013).

30. GBD Risk Factors Collaborators. Global, regional, and national comparative risk assessment of 79 behavioural, environmental and occupational, and metabolic risks or clusters of risks, 1990-2015: a systematic analysis for the Global Burden of Disease Study 2015. Lancet 388, 1659–1724 (2016).

31. Blood Pressure Lowering Treatment Trialists, C. et al. Blood pressure-lowering treatment based on cardiovascular risk: a meta-analysis of individual patient data. Lancet 384, 591–8 (2014).

32. Nakao, E. et al. Elevated Plasma Transforming Growth Factor beta1 Levels Predict the Development of Hypertension in Normotensives: The 14-Year Follow-Up Study. Am J Hypertens 30, 808–814 (2017).

33. Feng, W., Dell'Italia, L.J. & Sanders, P.W. Novel Paradigms of Salt and Hypertension. J Am Soc Nephrol 28, 1362–1369 (2017).

34. International PPH Consortium et al. Heterozygous germline mutations in BMPR2, encoding a TGF-beta receptor, cause familial primary pulmonary hypertension. Nat Genet 26, 81–4 (2000).

35. Voight, B.F. et al. Twelve type 2 diabetes susceptibility loci identified through large-scale association analysis. Nat Genet 42, 579–89 (2010).

36. Rossi, G.P. et al. A prospective study of the prevalence of primary aldosteronism in 1,125 hypertensive patients. J Am Coll Cardiol 48, 2293–300 (2006).

37. Calhoun, D.A., Nishizaka, M.K., Zaman, M.A., Thakkar, R.B. & Weissmann, P. Hyperaldosteronism among black and white subjects with resistant hypertension. Hypertension 40, 892–6 (2002).

38. Drelon, C., Berthon, A., Mathieu, M., Martinez, A. & Val, P. Adrenal cortex tissue homeostasis and zonation: A WNT perspective. Mol Cell Endocrinol 408, 156–64 (2015).

39. El Wakil, A. & Lalli, E. The Wnt/beta-catenin pathway in adrenocortical development and cancer. Mol Cell Endocrinol 332, 32–7 (2011).

40. Teo, A.E. et al. Pregnancy, Primary Aldosteronism, and Adrenal CTNNB1 Mutations. N Engl J Med 373, 1429–36 (2015).

41. Tissier, F. et al. Mutations of beta-catenin in adrenocortical tumors: activation of the Wnt signaling pathway is a frequent event in both benign and malignant adrenocortical tumors. Cancer Res 65, 7622–7 (2005).

42. Oliveira-Paula, G.H. et al. Polymorphisms in VEGFA gene affect the antihypertensive responses to enalapril. Eur J Clin Pharmacol71, 949–57 (2015).

43. Yang, R. et al. Hypertension and endothelial dysfunction in apolipoprotein E knockout mice. Arterioscler Thromb Vasc Biol 19, 2762–8 (1999).

44. Sofat, R. et al. Circulating Apolipoprotein E Concentration and Cardiovascular Disease Risk: Meta-analysis of Results from Three Studies. PLoS Med 13, e1002146 (2016).

45. Conrad, K.P. Unveiling the vasodilatory actions and mechanisms of relaxin. Hypertension 56, 2–9 (2010).

46. Sun, H.J. et al. Relaxin in paraventricular nucleus contributes to sympathetic overdrive and hypertension via PI3K-Akt pathway. Neuropharmacology 103, 247–56 (2016).

47. Pawlak, J.B., Wetzel-Strong, S.E., Dunn, M.K. & Caron, K.M. Cardiovascular effects of exogenous adrenomedullin and CGRP in Ramp and Calcrl deficient mice. Peptides 88, 1–7 (2017).

48. Miyamoto, Y. et al. Phosphatidylinositol 3-kinase inhibition induces vasodilator effect of sevoflurane via reduction of Rho kinase activity. Life Sci 177, 20–26 (2017).

49. Ohtsu, H. et al. Signal-crosstalk between Rho/ROCK and c-Jun NH2-terminal kinase mediates migration of vascular smooth muscle cells stimulated by angiotensin II. Arterioscler Thromb Vasc Biol25, 1831–6 (2005).

## References

50. Wain, L.V. et al. Novel insights into the genetics of smoking behaviour, lung function, and chronic obstructive pulmonary disease (UK BiLEVE): a genetic association study in UK Biobank. Lancet Respir Med 3, 769–81 (2015).

51. Bycroft, C.F., C; Petkova, D; Band, G; Elliot, LT; Sharp, K; Motyer, A; Vukcevic, D; Delaneau, O; O'Conell, J; Cortes, A; Welsh, S; McVean, G; Leslie, S; Donelly, P; Marchini, J. Genome-wide geentic data on 500,000 UK Biobank Participants. bioRxiv 166298 (2017).

52. Tobin, M.D., Sheehan, N.A., Scurrah, K.J. & Burton, P.R. Adjusting for treatment effects in studies of quantitative traits: antihypertensive therapy and systolic blood pressure. Stat Med 24, 2911–35 (2005).

53. Marouli, E. et al. Rare and low-frequency coding variants alter human adult height. Nature 542, 186–190 (2017).

54. Wain, L.V. et al. Genome-wide association study identifies six new loci influencing pulse pressure and mean arterial pressure. Nat Genet43, 1005–11 (2011).

55. Genomes Project, C. et al. A global reference for human genetic variation. Nature 526, 68–74 (2015).

56. Willer, C.J., Li, Y. & Abecasis, G.R. METAL: fast and efficient meta-analysis of genomewide association scans. Bioinformatics 26, 2190–1 (2010).

57. Purcell, S. et al. PLINK: a tool set for whole-genome association and population-based linkage analyses. Am J Hum Genet 81, 559–75 (2007).

58. Zhou, J. & Troyanskaya, O.G. Predicting effects of noncoding variants with deep learning-based sequence model. Nat Methods 12, 931–4 (2015).

59. Pers, T.H. et al. Biological interpretation of genome-wide association studies using predicted gene functions. Nat Commun 6, 5890 (2015).

60. Dunham, I.K., E.; Iotchkova, V.; Morganella, S.; Birney, E. FORGE: A tool to discover cell specific enrichments of GWAS associated SNPs in regulatory regions. F1000Research 4 (2015).

61. Wang, J., Vasaikar, S., Shi, Z., Greer, M. & Zhang, B. WebGestalt 2017: a more comprehensive, powerful, flexible and interactive gene set enrichment analysis toolkit. Nucleic Acids Res (2017).

62. Canela-Xandri. O.R., Konrad; Tenesa, Albert. An atlas of genetic associations in UK Biobank. bioRxiv 176834 (2017).

